# Genome-wide association study identifies 143 loci associated with 25 hydroxyvitamin D concentration

**DOI:** 10.1101/860767

**Authors:** Joana A Revez, Tian Lin, Zhen Qiao, Angli Xue, Yan Holtz, Zhihong Zhu, Jian Zeng, Huanwei Wang, Julia Sidorenko, Kathryn E Kemper, Anna AE Vinkhuyzen, Julanne Frater, Darryl Eyles, Thomas HJ Burne, Brittany Mitchell, Nicholas G Martin, Gu Zhu, Peter M Visscher, Jian Yang, Naomi R Wray, John J McGrath

**Author notes:** Equal contribution (Naomi R Wray, John J McGrath). Addresses for correspondence: Naomi Wray, John McGrath.

## Abstract

Vitamin D deficiency is a candidate risk factor for a range of adverse health outcomes. In a genome-wide association study of 25 hydroxyvitamin D (25OHD) concentration in 417,580 Europeans we identified 143 independent loci in 112 1-Mb regions providing new insights into the physiology of vitamin D and implicating genes involved in (a) lipid and lipoprotein metabolism, (b) dermal tissue properties, and (c) the sulphonation and glucuronidation of 25OHD. Mendelian randomization models found no robust evidence that 25OHD concentration had causal effects on candidate phenotypes (e.g. BMI, psychiatric disorders), but many phenotypes had (direct or indirect) causal effects on 25OHD concentration, clarifying the relationship between 25OHD status and health.

## Introduction

In recent decades there has been considerable interest in the links between vitamin D levels and general health. While classically linked to bone disorders, there is growing evidence to suggest that suboptimal vitamin D status may be a risk factor for a much wider range of adverse health outcomes^1^. Vitamin D, the ‘sunshine hormone’, is the precursor of a seco-steroid transcription regulator that operates via a nuclear receptor, and like other steroid hormones, exerts transcriptional control over many regions of the genome across many different tissues. In environments with access to adequate sunshine, ultraviolet radiation on the skin converts a precursor of cholesterol to vitamin D_3_. This is then further converted to 25 hydroxyvitamin D_3_ (25OHD; used in assays of general vitamin D status), and then to the active hormone 1,25 dihydroxyvitamin D_3_ (1,25OHD) in a variety of tissues. Some foods and vitamin D supplements also contribute to vitamin D levels. Definitions of vitamin D deficiency (e.g. < 25 nmol/L of 25OHD) are predominantly based on bone health^2^ – according to these definitions, vitamin D deficiency is common in many countries, regardless of latitude and economic status^3^.

Environmental factors such as season of testing and latitude contribute substantially to the serum concentration of 25OHD (lower in winter/spring; lower at higher latitudes)^4,5^. With respect to the genetic architecture of 25OHD, twin and family studies have reported a wide range of heritability estimates (from 0%^6^ to 90%^7^). A recent multivariate twin study demonstrated that approximately half of the total additive genetic variation in 25OHD may reflect genetic variation in skin colour and sun exposure behaviour^8^. Genome-wide association studies (GWAS) have identified common single nucleotide polymorphisms (SNPs) located in biologically plausible genes^9^. The largest GWAS to date (N = 79,366) reported six significant loci, which include *GC* (the vitamin D binding protein gene), the *DHCR7/NADSYN1* region (*DHCR7* is involved in a conversion of a 25OHD precursor molecule to cholesterol) and *CYP2R1* and *CYP24A1* genes (which encode enzymes involved in 25OHD metabolism^10^). In total, common SNPs explain 7.5% (standard error (s.e.) 1.9%) of the variance of 25OHD^10^.

Here, we conduct a GWAS of 25OHD based on the large UK Biobank (UKB) sample^11^ and conduct a suite of post-GWAS analyses to aid interpretation of the results (**Figure 1**). We present models that explore the genetic/causal relationship between body mass index (BMI) and 25OHD (high BMI is associated with lower 25OHD concentration in observational studies)^12^. Because we have an interest in the association between 25OHD and mental disorders^13^, we use Mendelian randomization methods to investigate the bidirectional association between 25OHD and psychiatric disorders, as well as with a wider range of traits and diseases. Additionally, we present a GWAS to identify loci associated with variance in 25OHD (i.e., variance quantitative trait locus (vQTL) analysis) which can identify putative genotype environment interactions without prior identification of the environmental effect^14^.

**Figure 1:**
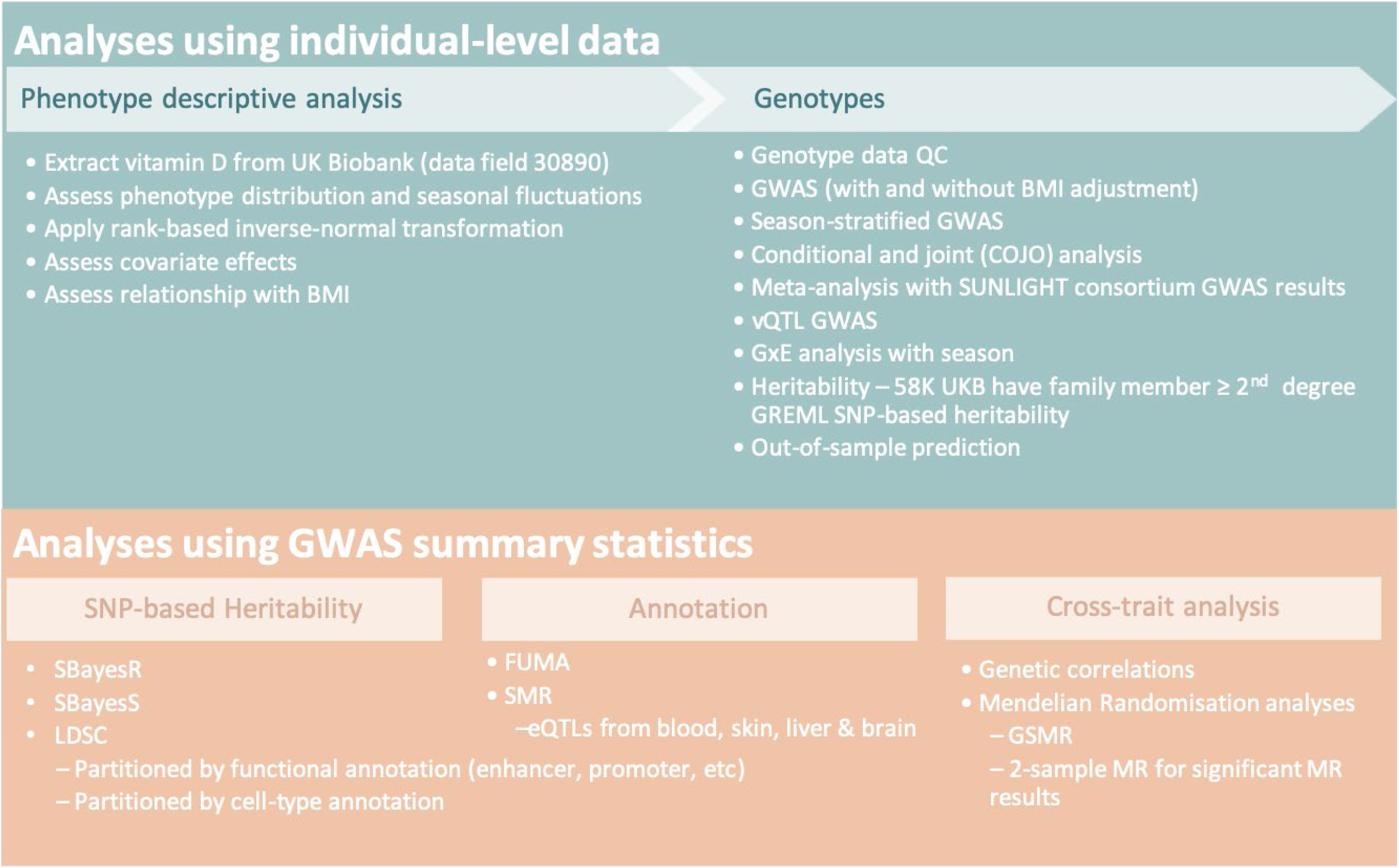
Outline of key analytic steps described in this study. Abbreviations: BMI, body mass index; eQTL, expression quantitative trait locus; FUMA, functional mapping and annotation^27^; GREML^79^, genomic relationship restricted maximum likelihood^78^; GSMR, generalized summary-based MR; GWAS, genome-wide association study; GxE, genotype by environment interaction; LDSC, linkage disequilibrium score regression; MR, Mendelian Randomisation; SMR, summary-based MR^30^; QC, quality control; UKB, UK Biobank; vQTL, variance quantitative trait locus^14^.

## Results

417,580 European UKB participants had both measures of vitamin D 25OHD and genome-wide genotypes (**Methods**). The distribution of 25OHD concentration, in keeping with expectation, is right skewed (**Supplementary Figure 1a**), and showed the expected seasonal fluctuation (**Supplementary Figures 1b and 1e**), with median, mean and interquartile range of 47.9, 49.6, 33.5 – 63.2 nmol/L (**Supplementary Table 1**). Covariates of age, BMI, genotyping batch, assessment centre, month of testing, supplement intake and the first four ancestry principal components (PCs), but not sex, were all significantly associated with 25OHD (**Supplementary Table 1**). Month of testing accounts for 14% of the variance of 25OHD. Subsequent analyses use 25OHD after rank-based inverse-normal transformation (RINT) unless otherwise stated.

### Heritability and SNP-based heritability

Our UKB sample included a set of 58,738 individuals related with coefficient of relationship (*r*) > 0.2 to at least one other person in the set (“all relatives”), from whom we estimate the heritability of 25OHD to be 0.32 (s.e. = 0.01) with little evidence for inflation from shared family environment (**Figure 2, Supplementary Figure 2, Supplementary Table 2**). The SNP-based heritability estimate 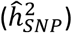, which captures the genetic contribution from common (minor allele frequency or MAF > 0.01) variants, was 0.13 (s.e. = 0.01) (see **Supplementary Figure 2, Supplementary Table 2** for a comparison of 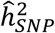 estimated from various methods). 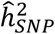 was significantly higher (*P* = 1.5 × 10^−3^) when estimated only from individuals measured for 25OHD in summer months (June to October) compared to those measured in winter months (December to April) (0.19, s.e. = 0.02 *vs.* 0.10, s.e. = 0.02) (**Figure 2**), as found for estimates of twin heritability^8^. The genetic correlation between the seasons was 0.80 (s.e. = 0.11), not significantly different from 1. The proportion of SNPs estimated to have an effect on the trait (polygenicity parameter) using the SBayesS method^15^ was 0.8% or 9,000 SNPs of the ∼1.1 million HapMap3 panel^16^ common SNPs (**Supplementary Table 3**), much lower than estimates for most complex traits^15^. The SBayesS *S* parameter, which describes the effect size-MAF relationship, was estimated as −0.78 (s.e. = 0.04; **Supplementary Table 4**), consistent with a model of negative selection on the genetic variants associated with 25OHD levels (the magnitude of *S* is higher than those of most complex traits studied^15^). Estimation of 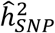 partitioned into 10 components based on five MAF bins (each median-split by linkage disequilibrium score) did not provide strong evidence for an increased role for less common variants, given the s.e. of estimates (**Supplementary Figure 3**). Despite a strong phenotypic association between 25OHD and BMI of −0.76 nmol/L/BMI unit (−0.036 RINT(25OHD) standard deviation (SD) units/BMI unit, *P* < 2.2 × 10^−16^) and a phenotypic correlation of −0.17 (**Supplementary Table 1**), the estimates of heritability (both family and SNP-based) were hardly impacted when BMI was included as a covariate (**Supplementary Figure 2**).

**Figure 2.**
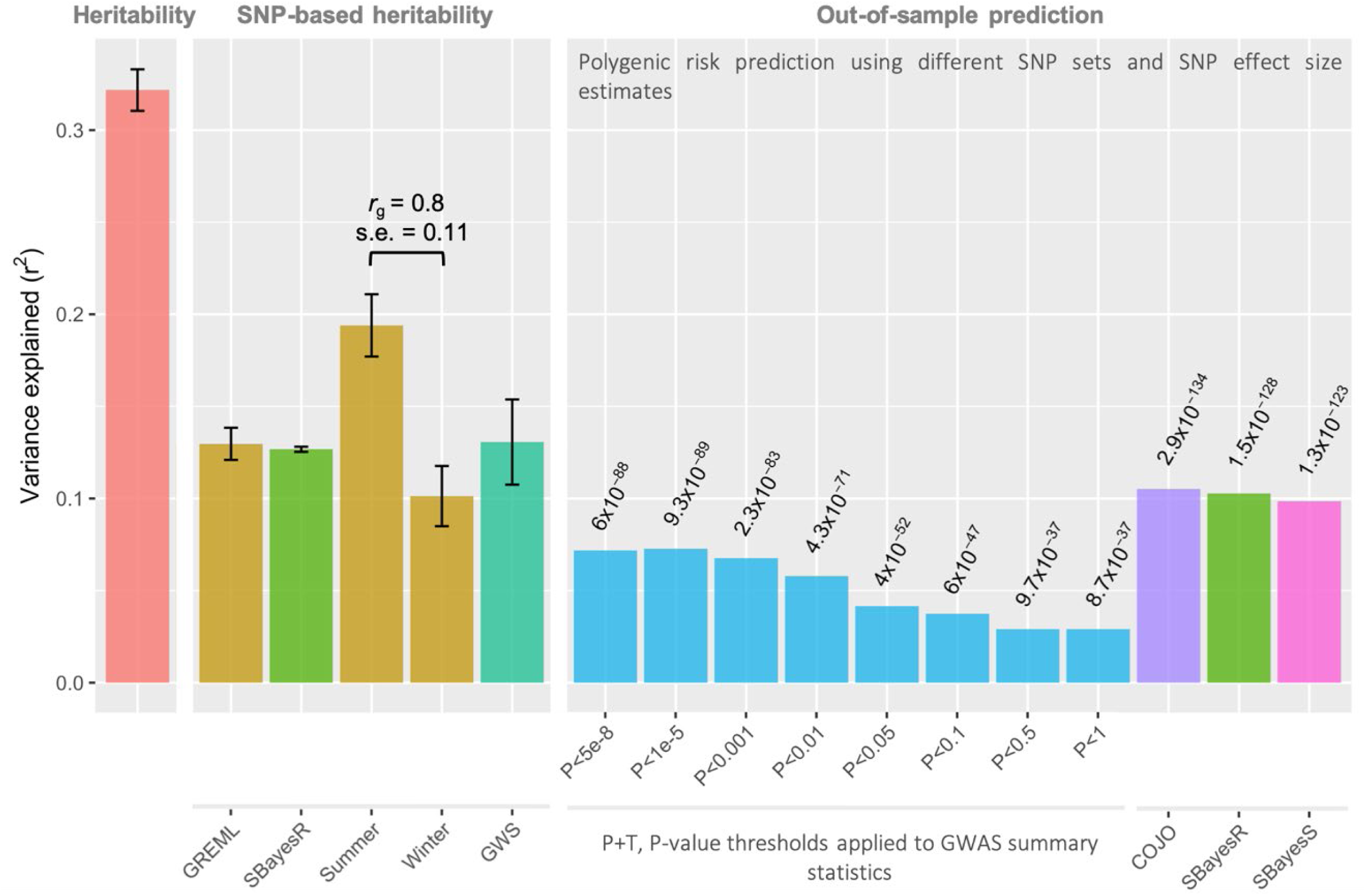
Heritability and SNP-based heritability estimates and variance explained in out-of-sample prediction. GCTA-GREML was used to estimate heritability (N=58,738 relatives) and SNP-based heritabilities labelled GREML (GREML), summer, winter for samples of ∼50K randomly drawn from the UKB. The SBayesR SNP-based heritability is estimate from the GWAS summary statistics (N=417,580). The GWS estimate used only genome-wide significant COJO loci identified from the UKB GWAS in a GCTA-GREML analysis using the QIMR sample (N = 1,632). In out-of-sample prediction into the QIMR sample, polygenic scores calculated by the standard *P*-value threshold method was outperformed by calculating the score from COJO identified GWS SNPs, and by using SNP effect estimates calculated from GWAS summary statistics using the SBayesR or SBayesS methods. Abbreviations: COJO, conditional and joint; GWS, genome-wide significant; r_g_, genetic correlation; s.e., standard error.

### Genome-wide association study (GWAS) analysis

Given the potential for collider bias from using a heritable trait as a covariate^17^, we conducted GWAS for 25OHD with and without BMI as a covariate. We also used mtCOJO^18^ to estimate the 25OHD SNP effects conditioning on those estimated for BMI from UKB data^19^, a summary-data-based conditional-analysis approach that was shown in simulations to be robust to collider bias when conditioning on a correlated trait^18^. Results were comparable across the three levels of BMI adjustment (**Supplementary Table 5)**, so we report those with no correction for BMI, using results from all three analyses when this aids interpretation of results.

A total of 8,806,780 SNPs with MAF > 0.01 were tested in the GWAS analysis. Of these, 18,864 were genome-wide significant (GWS; *P* < 5 × 10^−8^). To identify independently associated loci, we applied the GCTA-COJO method^20^ to the GWAS summary statistics using LD between SNPs estimated from a UKB subset (**Methods**), and identified 143 independent loci (including one on chromosome X) (**Figure 3; Supplementary Table 6**) in 112 1-Mb regions. Of these, 15 loci were low frequency variants (MAF < 0.05), and 106 regions had no previously identified associations. All six loci reported in previous vitamin D GWAS^10,21,22^ were replicated in our study. While recognising that the COJO method cannot distinguish between SNPs in perfect LD, we note that within the 143 COJO independent variants: (a) 14 were non-synonymous variants that alter protein coding (*NRIP1, DSG1, TM6SF2, PLA2G3, GCKR, APOE, PCSK9, SEC23A, FLG, NPHS1, SDR42E1, CPS1, ADH1B, UGT1A5*), and (b) 9 were annotated to include small insertion/deletions. A summary of results is provided in **Figure 4**, but are discussed later.

**Figure 3:**
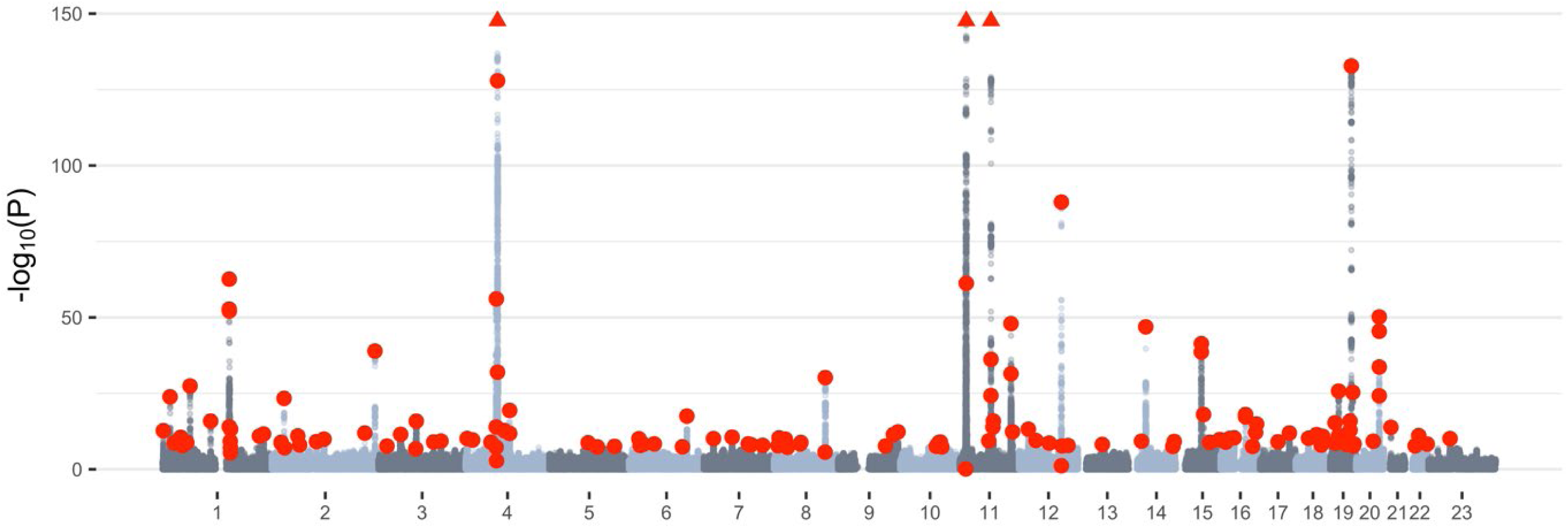
Manhattan plot of the 25OHD GWAS in the UK Biobank. Manhattan plot showing the −log_10_ *P-*values from association of 25 hydroxyvitamin D (25OHD). Red dots represent independent variants identified as genome-wide significant with conditional and joint analysis (COJO)^20^ applied to the GWAS summary statisitics. The horizontal axis shows each chromosome, with 23 representing the X chromosome. The vertical axis is restricted to −log_10_ *P*-values < 150. Five COJO SNPs with *P* < 1×10^150^ (four on chromosome 11 and one on chromosome 4; Supplementary Table 6) with approximate locations represented by three red triangles at the top edge of the plot.

**Figure 4:**
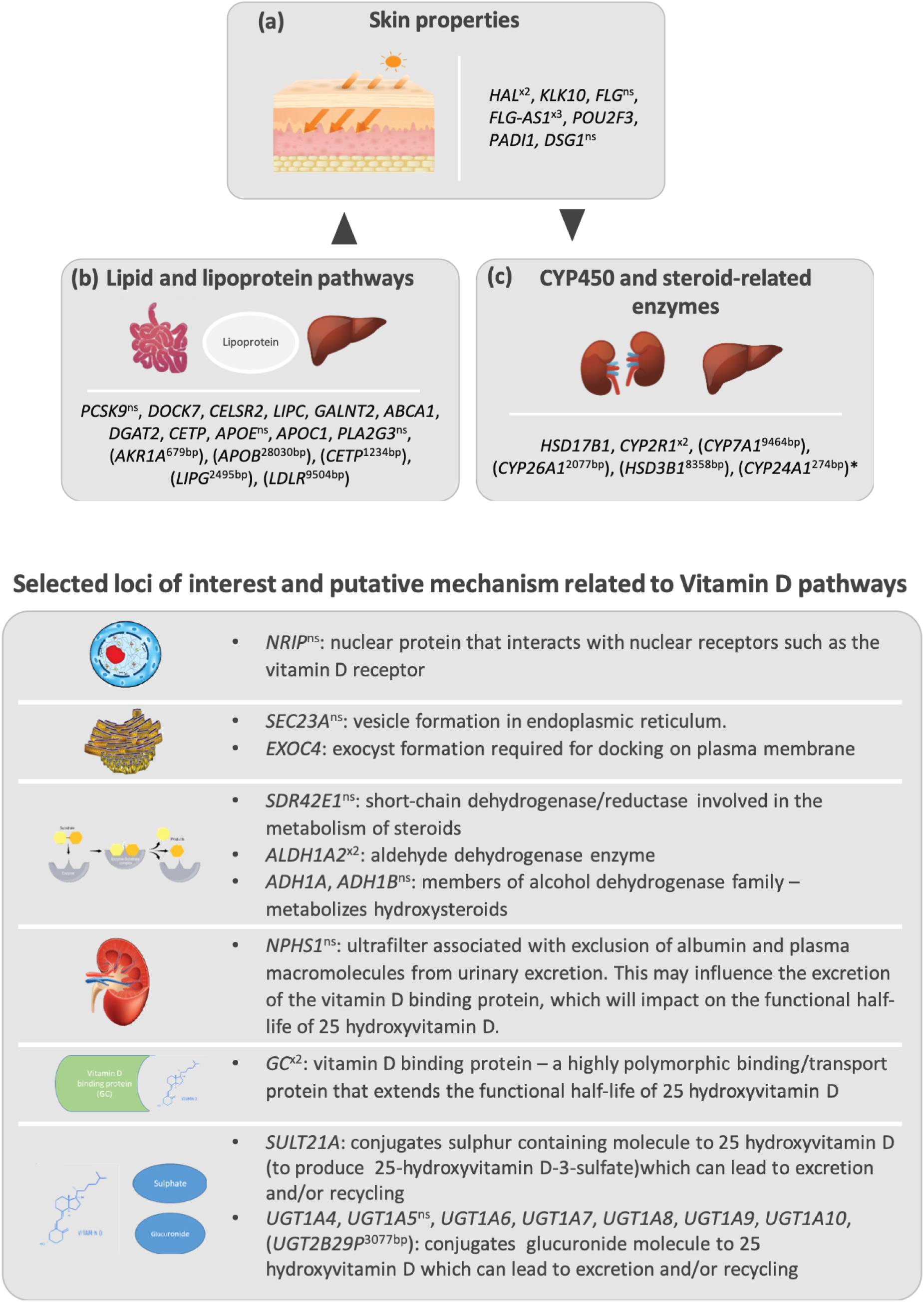
Summary of selected variants associated with 25 hydroxyvitamin D in the UK Biobank. Top panel shows loci associated with skin integrity, lipid and lipoprotein pathways, and CYP450 and steroid associated enzymes. Lower panel shows selected variants and putative mechanisms related to 25 hydroxyvitamin D concentration. For selected inter-genic loci, the nearest (upstream or downstream) gene is shown in brackets. The distance between the loci and the nearest gene is shown in base pairs. \**CYP24A1* was also the closest gene for an additional three inter-genic loci with distances between 32,865-55,282 base pairs. Abbreviations: ns, non-synonymous variant; x2 or x3, two or three loci found within the gene.

Summary statistics from the SUNLIGHT consortium^10^ were available for 2,579,297 SNPs and the genetic correlation estimate with UKB results was not significantly different from 1 (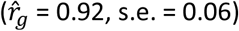. Meta-analysis with our UKB GWAS results after imputation^23,24^ of the SUNLIGHT summary statistics (**Supplementary Methods**) (6,912,294 overlapping SNPs) identified 15,154 GWS variants, 150 GCTA-COJO independent SNPs (**Supplementary Methods, Supplementary Table 5**). Given that the meta-analysis only increased the number of significant loci by seven, and given our preference not to include BMI as a covariate, we continued with the UKB-only results for our downstream analyses. It is notable, that random draws of ∼80K people from the UKB, the same size approximately as from the SUNLIGHT consortium meta-analysis, identified ∼20 independent COJO GWS loci (Supplementary Figure 4a), a 36% increase compared to 14 COJO SUNLIGHT consortium loci, demonstrating the power gained for equal sample size from having a single cohort study, as previously shown for height and BMI^25^. Here, we find an approximately linear relationship between sample size and GWS discovery of 3.7 loci/10K people (**Supplementary Figure 4b**).

### Replication and out-of-sample genetic risk prediction

To get an unbiased estimate of the phenotypic variability explained by the independent SNPs in our GWAS, we conducted a GREML analysis in the independent QIMR dataset^8^ (N = 6,233, N = 1,632 unrelated). UKB genome-wide significant COJO SNPs explained 13% of the variance in RINT(25OHD) residuals (i.e. after accounting for covariates) when fitted jointly. Polygenic prediction into the QIMR sample using SNP effects estimated in the UKB and the standard *P*-value thresholding method explained a maximum of 7.3% of the variance in RINT(25OHD)(**Figure 2; Supplementary Table 7**) (*P* = 9.3 × 10^−89^, at *P*-value threshold of *P* < 1 × 10^−5^). As expected, when the PRS were derived from SNP weights from COJO or Bayesian methods applied to the GWAS summary statistics^15,26^ the prediction variance was higher, to a maximum of 10% (**Figure 2; Supplementary Table 7).**

### Functional mapping and annotation of GWAS

To annotate the 25OHD GWAS, we first used the FUMA online pipeline^27^. Gene-set analyses showed that the top four pathways were related to glucuronidation, ascorbate and aldarate metabolism, and uronic acid metabolism (**Supplementary Tables 8 and 9**). Keratinization was the top Gene Ontology (GO) biological processes identified. Based on 53 tissue types from GTEx v6^28^ the top tissues for differentially expressed genes identified in the GWAS were liver, brain and skin (sun exposed, and non-sun exposed; **Supplementary Table 10**). Partitioned SNP-based heritability analysis^29^ using cell-type-specific annotations identified five cell types (hepatocytes, two types of liver cells, skin cells and blood cells) at the nominal significance level of 0.05 (**Supplementary Table 11**), but none remained significant after correction for multiple testing (*P* < 2.4 × 10^−4^). In partitioned SNP-based heritability analysis using SNP annotation to 53 functional categories^29^, 11 passed multiple testing significance threshold (*P* < 9.4 × 10^−4^; **Supplementary Table 12**) with a mix of annotations including transcription factor binding sites and transcription start sites (notable because vitamin D operates via a nuclear receptor, which binds to vitamin D response elements), as well as a role for repressed sites, conserved regions, enhancer and coding regions, and histone modification marks.

To identify 25OHD SNP associations with statistical evidence consistent with a causal/pleiotropic association via gene expression, we used summary-data-based Mendelian randomization (SMR)^30^ using the 15,504 gene probes with significant cis-eQTLs identified from whole blood eQTLGen data^31^. After Bonferroni correction, we found 112 significantly-associated gene expression probes (*P*_SMR_ < 3.2 ×10^−6^, i.e., 0.05 / *m*, with *m* = 15,504, being the total number of probes tested in SMR analysis; **Supplementary Table 13, Supplementary Figure 5; <u>Supplementary Data</u>**). These results are discussed in detail in the **Supplementary Note** and add weight to the hypothesis that the SMR identified eQTL variants may be causally related to 25OHD concentrations.

### Genetic correlations and putative causal relationships with other traits

First, we investigated the relationship between 25OHD and BMI. The LDSC^32^ genetic correlation estimated from 25OHD and BMI GWAS summary statistics was −0.17 (s.e. = 0.03) (**Supplementary Figure 2, Supplementary Table 14**). Bidirectional Mendelian randomisation^18^ analysis provided strong support for the hypothesis that high BMI is causal for low 25OHD (b_BMI.25OHD_ = −0.130; s.e. = 0.005; *P* = 4.7 × 10^−162^; based on 1,020 BMI-associated SNP instruments), with no support for a causal effect of vitamin D on BMI (b_25OHD.BMI_ = 0.008; s.e. = 0.006; *P* = 0.20; based on 210 vitamin D-associated SNPs) (these results were confirmed by other MR methods^33^; **Supplementary Table 15**). Notably, the HEIDI-outlier test in the GSMR analyses excluded 70 BMI and 67 25OHD SNP instruments, whose combination of SNP effect sizes likely reflects a pleiotropic relationship or confounding. Using the SNPs excluded by the HEIDI-outlier test, the estimates were b_BMI.25OHD_ = 0.17 (s.e. = 0.0182; *P* = 1.2 × 10^−20^) and b_25OHD.BMI_ = −0.15 (s.e. = 0.017, *P* = 2.7 × 10^−18^). Hence, despite the clear evidence for a causal relationship between high BMI and low 25OHD, the biological relationship between these traits is more complex.

Next, we estimated genetic correlations (*r*_*g*_) between 25OHD and 746 traits with GWAS summary statistics available in LD Hub^34^, and we used LDSC to estimate *r*_*g*_ between 25OHD and 18 traits (including six psychiatric disorders) with GWAS summary statistics that are more recent than those included in LD Hub. Although many of the traits are highly correlated, we use a Bonferroni correction for 764 tests as the threshold for discussion of *r*_*g*_. We found significant associations between 25OHD and a range of brain-related phenotypes (including autism spectrum disorder, intelligence, major depressive disorder, bipolar disorder and schizophrenia; **Supplementary Figure 6**). Notably, the most significant *r*_*g*_ were with cognitive-associated traits — for example, a negative correlation 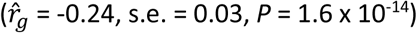 with intelligence. There was also a significant negative *r*_*g*_ with hours spent using a computer 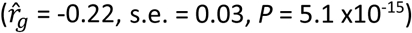. These findings may be mediated by an association between higher intelligence and behaviour associated with less exposure to bright sunshine (and thus, lower 25OHD). Of note, behaviours associated with outdoor activity (duration of walks, duration of vigorous activity) were positively associated with 25OHD, while phenotypes related to chronic disability were negatively associated with 25OHD.

Next, we investigated if some of the significant genetic correlations could be explained by causal relationships using bidirectional GSMR models – here a more complex pattern of association emerged (**Figure 5, Supplementary Table 16**). We found no evidence for putative causal effects between 25OHD and other traits; GSMR analyses without the HEIDI-outlier filtering step (**Figure 5a**) suggest strong pleiotropy for some traits like dyslipidemia, coronary artery disease, intelligence and educational attainment. Finally, we examined the reciprocal relationship – if variants associated with a range of traits were directionally associated with 25OHD. Regardless of the use of HEIDI filtering, and often regardless of adjustments for BMI, we found evidence consistent with increased risk of several traits or disorders being causal (directly or indirectly) with lower 25OHD concentrations. This was the case for intelligence, dyslipidemia, major depression, bipolar disorder, type 2 diabetes and schizophrenia. The findings might suggest these traits or disorder are associated with behaviours that lead to reduced production of 25OHD (e.g. less outdoor activity and physical activities). The GSMR findings were also checked with the portfolio of MR methods implemented in the 2-sample MR (2SMR) software^33^ (**Supplementary Table 17**).

**Figure 5:**
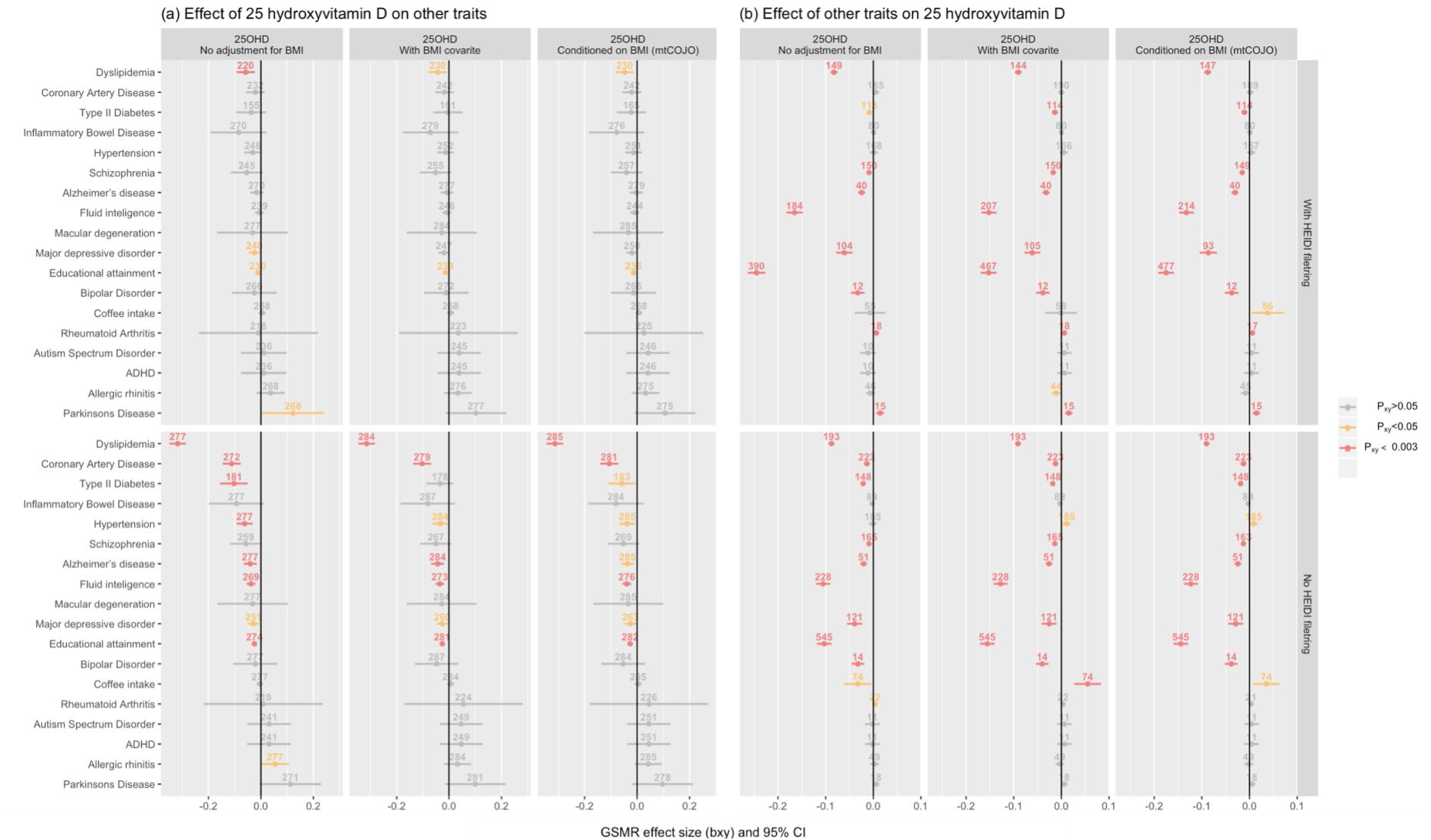
Bidirectional Generalized Summary data level Mendelian Randomization (GSMR) between 25 hydroxyvitamin D concentrations and selected phenotypes, by three types of adjustments for body mass index (BMI) and with/without HEIDI filtering of pleiotropic loci. Panel (a) shows the estimate of the causal effect (dots), and 95% confidence intervals (bars), of 25 hydroxyvitamin D concentration (25OHD) on selected phenotypes. Negative GSMR effect sizes indicate that variants associated with increased 25OHD concentration were associated with a smaller value/reduced risk for the phenotypes of interest. Panel (b) shows the estimate of the causal effect (and 95% confidence intervals) of the same selected phenotypes on 25OHD concentration. Results are presented with (upper half) and without (lower half) filtering pleiotropic associations with the heterogeneity in dependent instruments (HEIDI) test, respectively. The numbers above each effect size indicate the number of SNP instruments used in each analysis. For each set of analyses, we show GWAS results (i) without adjustment for body mass index (BMI), (ii) with BMI included as a covariate, and (iii) conditioned on BMI using mtCOJO^18^. The GMSR estimates and 95% confidence intervals are shown in three colours according to the *P-*value thresholds.

### Proxy-environment vQTL, season analysis and gene by environment interaction

We conducted a genome-wide vQTL analysis, as implemented in OSCA^14^ to identify SNPs associated with variance in 25OHD (not RINT transformed). Such associations can reflect genotype-by-environment interaction in the absence of measurement, or indeed knowledge, of the interacting environmental risk factor. Using data from 318,851 unrelated individuals of European ancestry, we tested 6,098,063 variants with MAF > 0.05, and identified 4,008 GWS vQTLs, of which 25 were independent (LD *r*^2^ < 0.01, 5-MB window), and several were in well-characterized genes (e.g. *GC, UGT2B7, SEC23A, SULT2A1, KLK10, NADSYN1*). Of the 25 independent vQTLs, 23 were also QTLs (identified as genome-wide significant in the GWAS analysis) while the two non-QTL loci were still associated at *P*_GWAS_ < 10^−5^ (**Supplementary Table 18**). One was in the *POR* gene, which encodes a cytochrome p450 oxidoreductase that donates electrons from NADPH to cytochrome P450 enzymes (encoded by *CYP450* genes), which are involved in vitamin D metabolism^35^. Variants in *POR* have previously been associated with coffee intake^36^. The other exclusive vQTL (rs1030431) is 12,126 bp upstream from *UBXN2B*; the SNP is significantly associated with gall bladder diseases and lipid metabolism traits in the UKB^37^.

An environmental factor with known association with 25OHD is the season of testing. To investigate if the associations between the vQTLs and the phenotypic variance of 25OHD reflected gene-environment (GxE) interactions with season of blood draw, we performed a GxE analysis with season (winter vs. summer). Of 6,098,063 variants tested (MAF > 0.05), 1,127 had a GWS (*P* < 5 × 10^−8^) interaction with season, and 1,120 (99%) were also GWS in the vQTL analysis. From the 1,127 GWS interactions, five were independent (LD *r*^2^ < 0.01, window 5 Mb) and were located in regions that have well-known vitamin D related genes in chromosomes 7, 11, and 14 (**Supplementary Table 19)**. Notably, of the 20 vQTL loci without significant GxE with season, at least half showed no evidence at all for GxE with season (**Supplementary Figure 7**), so these variants are candidates for GxE with other environmental factors.

## Discussion

We have identified 143 loci associated with 25OHD concentration. Recognising that only six associated loci had been reported to date these discoveries provide important new insights into previously unknown or poorly understood vitamin D-related pathways and substantially increase our knowledge of the genetic correlates of 25OHD compared to previous studies^9^ (**Figure 4**). First, the three most associated loci, all identified in previous studies^10^, are noteworthy (chr4:rs1352846, chr11:rs116970203 and chr11:rs12794714, all *P* < 1.0 × 10^−400^, all with their minor allele reducing 25OHD). rs1352846 (MAF = 0.29 (G)) is in the *GC* locus^10,22^, which encodes a protein synthesized in the liver that binds to, and transports vitamin D and its metabolites. rs116970203 is a low frequency variant (MAF = 0.03 (A)) located in intron 11 of the *PDE3B* gene. It is also a perfect proxy for rs117913124 (LD *r*^*2*^ = 1), a low frequency synonymous coding variant in *CYP2R1*, which was previously reported to associate with 25OHD^21^. Another *CYP2R1* synonymous variant was also identified (rs12794714; MAF = 0.42 (A)). *CYP2R1* encodes a crucial hepatic enzyme involved in the hydroxylation of vitamin D to 25OHD. Given the complexity of the association pattern observed in chromosome 11, we confirmed the independence of the COJO identified variants using individual-level data (**Supplementary Table 20**). In line with previous findings^38^, the two-way conditional analysis showed that the effect of the low frequency SNP (rs116970203 or rs117913124) and common SNP (rs12794714 or rs10741657), were largely independent.

Our findings provide convergent evidence that genes related to lipid- and lipoprotein-related pathways influence 25OHD concentration. In particular, we confirm a unidirectional relationship between SNP instruments that influence higher BMI and lower 25OHD concentration, but not the reciprocal relationship. This relationship exists against a background of a highly intercorrelated pattern of relationships between genes that influence both 25OHD and a wide range of lipid-related metabolic phenotypes. There were variants within genes with well-described functions related to lipid and lipoprotein related pathways^39^ (e.g. *PCSK9, DOCK7, CELSR2, GALNT2, ABCA1, DGAT2, CETP, APOE, APOC1, PLA2G3*). In addition, several inter-genic loci had ‘closest’ upstream or downstream genes of interest to lipid and lipoprotein pathways (*AKR1A, APOB, CETP, LIPG, LDLR*). Variants in these genes influence overall lipid concentrations, including the concentration of 7-dehydrocholesterol in the skin. We identified a locus (chr11:rs12803256) in an uncharacterized RNA gene (*FLJ42102*) 11,057 base pairs upstream from *DHCR7*. This region has been identified in previous GWAS studies, and *DHCR7* is a strong candidate gene because of its known role in the conversion of 7-dehydrocholesterol in the skin to pre-vitamin D_3_. We note that the broad region on Chr 11 containing *DHCR7* and *NADSYN1* included several loci of interest according to both GCTA-COJO and SMR analyses – this complex area warrants additional research.

The GWAS uncovered a range of novel findings indicating that properties of the skin not related to pigmentation are associated with 25OHD concentration. While it is well known that individuals with darker skin tend to have lower 25OHD (related to the melanin content in the skin blocking UVB)^1,8^, our findings provide evidence that SNPs associated with genes that influence dermal development (e.g. *PADI*)^40^ and integrity (e.g. *FLG*; *FLG-AS1, POU2F3, KLK10, DSG1*)^41-44^ are also associated with 25OHD status. It has been suggested that variants in the *FLG* gene may have evolved in order to optimize 25OHD production at high latitude^45,46^. *HAL* (histidine ammonia-lyase) codes for an enzyme that deaminates L-histidine to trans-uronic acid. The top SNP in this region (rs10859995) is within an intron of this gene. The gene is expressed in the skin, and is upregulated during keratinocyte differentiation^47^. It has been demonstrated that trans-urocanic acid in the stratum corneum can absorb UVB^48^ and can reduce the production 25OHD^49^. The MAGMA gene-set analysis^27^ also showed that variants associated the uronic acid pathways were significantly over-represented in our findings (**Supplementary Table 9**). The concentration of trans-uronic acid varies widely between individuals^49,50^ but is not related to skin colour/pigmentation^50^. It is important to note that our sample was restricted to Europeans and analyses included ancestry PCs as covariates, four of which were strongly associated with 25OHD (**Supplementary Table 1**). If these PCs capture variants related to skin colour within Europeans, these variants are less likely to be identified in our analyses. FUMA analyses did not identify an over-representation of variants known to be related to skin colour in our GWAS.

Our study expands the range of enzymes implicated in the synthesis and breakdown of vitamin D related molecules. These include genes from the hydroxysteroid 17-beta dehydrogenase family (*HSD17B1, HSD3B1*), a family of short-chain dehydrogenases/reductases, which are involved in steroidogenesis and steroid metabolism^51^. *CYP2R1* is a key regulator of 25OHD status, via hepatic conversion of vitamin D to 25OHD — two loci were found within this gene. Other members of this large family of enzymes associated with 25OHD concentrations include *CYP7A1, CYP26A1*, and *CYP24A1*.

We identified many variants within genes related to the modification of lipophilic molecules (including seco-steroids such as 25OHD and related species). Associated regions on chromosomes 2 and 4 include enzymes in the UDP-glucuronosyltransferase family, which are critical in the glucuronidation pathways. The involvement of these genes in the degradation and potential conjugate recycling of 25OHD has recently been described^52,53^. We identified variants in the *SULT21A* gene, which encodes the enzyme responsible for the sulphonation of 25OHD^53,54^. Our findings provide support for the hypothesis that these mechanisms influence 25OHD concentration. We identified variants in the *SLCO1B1* gene, which encodes a transmembrane receptor that mediates the sodium-independent uptake of numerous endogenous compounds, including sulphated steroid molecules^55^. It is not known if this mechanism is involved in the uptake of the sulphated 25OHD, but metabolic studies have identified associations between variants in the *SLCO1B1* gene and a wide range of small molecules^56^. It has been proposed that vitamin D may undergo conjugate cycling (e.g. bidirectional conversion between 25OHD and 25OHD-sulphate)^57^. A proportion of total 25OHD may exist in the sulphated form, which could act as circulating reservoir for later de-sulphation in peripheral tissues. In addition, conjugated versions of 25OHD with glucuronide^52^ and sulfate^53^ have both been detected in bile, which suggests enterohepatic mechanisms may provide another reservoir that buffers total 25OHD reserves. The findings also have implications for how to assay total 25OHD reserves. Current extraction and assay techniques used to quantitate 25OHD are not optimized for sulphonated or glucuronidated species of 25OHD^58^, thus total 25OHD status may not accurately reflect the contribution of these conjugated species. In addition, these mechanisms would contribute to the functional half-life of 25OHD, and thus influence vitamin D status during periods of reduced exposure to bright sunshine (e.g. during winter). Finally, variants in a range of novel enzymatic pathways were also associated with 25OHD concentration (e.g. short-chain dehydrogenase/reductase, aldehyde dehydrogenase, alcohol dehydrogenase).

The large sample size afforded by the UKB sample, provides for the first time a description of the genetic architecture of 25OHD. The 143 loci explain 13% of the variance in an independent sample^8^, when fitted jointly (equalling the SNP-based heritability) and 10% of variance using a polygenic score predictor using SNP effect sizes estimated in the UKB. Since the latter is achieved from considering only genome-wide significant loci it means that a large part of the common variant signal is already well-captured and estimated by our sample size. In total, all genotyped/imputed variants with MAF > 0.01 explain about 41% of the heritability estimated from close relatives (heritability 0.32, SNP-based heritability 0.13, **Figure 2**). We estimate that about 9,000 common SNPs affect variation in 25OHD, and report evidence of negative selection through the SBayesS *S* parameter of −0.78 (which represents the relationship between MAF and effect size, and which is zero under a neutral model). The 143 loci represent only 112 1-Mb regions, with six of the 1-Mb regions harbouring four loci each. The final set of 143 loci was achieved by applying the COJO (conditional and joint) algorithm onto the GWAS summary statistics using the linkage disequilibrium structure to account for the correlation structure between SNPs. Two regions on chromosome 11 are particularly complex with the maximum change in significance observed for SNP rs61883501 (*P* = 0.749, *P*_COJO_ 4.0×10^−9^ and with a change in direction of effect, **Supplementary Table 6**). An increase in out-of-sample prediction from 7.3% from the standard *P*-value thresholding method to 10.5% when using COJO SNP effect estimates provides independent support for the validity of the approach.

We also identified 25 independent SNPs associated with variance in 25OHD — these are putative GxE loci. While 5 of these have strong evidence of interacting with season of measurement, at least 10 are GxE candidates with yet-to-be-identified environmental risk factors, and search of published GWAS results for association with these SNPs (i.e., PheWAS^37^) may help with this prioritization (**Supplementary Table 18**). In summer months the mean 25OHD concentrations are higher and a larger proportion of the variance could be attributed to genetic factors in summer compared to winter (SNP-based heritability of 0.19, s.e. = 0.02, vs 0.10, s.e. = 0.02, P_different_=1.5 × 10^−3^). Five loci were identified as significant in GxE analysis with season, and for two the direction of effect was reversed (**Supplementary Table 19**). The vitamin D phenotype is an interesting one to explore from the perspective of GxE as seasonal fluctuations provide a natural experiment to dissect components of the genetic architecture that influence synthesis (i.e. inflow) and excretion (i.e. outflow) of 25OHD-related pathways.

In the UKB participants, high BMI is associated with reduced 25OHD concentration, in keeping with a large body of observational epidemiology^59^. However, we did not find statistical evidence in support of a causal role for 25OHD level on BMI. In contrast, there was evidence for pleiotropic effects of SNPs on the two traits as well as for high BMI being causal (directly or indirectly) for low 25OHD. Genetic correlations were significant between 25OHD concentration and a range of phenotypes (**Figure 5**). However, in robust directional models, we found no evidence in support of a causal role for 25OHD concentration on these traits. Of interest, we found evidence that higher intelligence and an increased risk of several psychiatric disorders may cause reduced 25OHD concentrations. With respect to intelligence, this would be consistent with previous links between intelligence and years of education leading to working indoors, and subsequent lower concentrations of 25OHD^60,61^. One of our motivations for undertaking this study was to investigate the hypothesis of a causality relationship between 25OHD and psychiatric disorders^62^. The Mendelian randomization analyses conducted here do not support a causal role for 25OHD levels and these disorders, and hence the reported epidemiological associations could reflect confounding and/or reverse causation. Vitamin D deficiency is common in those with established psychiatric disorders, as a consequence of reduced outdoor behaviour^63,64^. It is feasible that the observed association between 25OHD concentration in blood spot samples taken at birth with later-life increased risk of schizophrenia^13,65^ could be confounded by outdoor behaviour of mothers, which may be correlated with the mother’s genetic liability to schizophrenia. While we find no evidence to support the hypotheses that variants associated with low 25OHD concentrations were associated with any of the selected phenotypes, we note that there is a linearity assumption in our Mendelian randomization analyses. In other words, if only very low concentrations of 25OHD are associated with adverse outcomes, then this non-linear exposure-risk association may not be confidently detected.

## Conclusions

We have identified 143 loci associated with 25OHD concentration, and have provided new directions for vitamin D research. In particular, our findings suggest that pathways related to sulphonation and glucuronidation warrant closer scrutiny – for example, there may be a case to measure these modified species of 25OHD and related molecules in order to better understand vitamin D status. Our studies based on Mendelian randomization do not support hypotheses that vitamin D concentration is associated with a broad range of candidate phenotypes, in particular, psychiatric disorders. The findings provide new insights into the physiology of vitamin D and the relationship between 25OHD status and health.

## Acknowledgements

This current study was carried out under the generic approval from the NHS National Research Ethics Service and conducted using the UK Biobank resource under projects 12505 and 10214. We thank the UKB participants, project team and funders for providing this important research resource. We thank the eQTLGen consortium for providing the cis eQTL dataset based on N = 32K participants. The Genotype-Tissue Expression (GTEx) Project was supported by the Common Fund of the Office of the Director of the National Institutes of Health, and by NCI, NHGRI, NHLBI, NIDA, NIMH, and NINDS. Funding for the QIMR sample was provided by the Australian National Health and Medical Research Council (NHMRC) and further supported by NHMRC Project Grants (1007677, 1099709) and a John Cade Fellowship (1056929). NHMRC also support Naomi Wray (1113400, 1078901), Peter Visscher (1113400, 1078037) and Jian Yang (1113400). Jian Yang is supported by the Australian Research Council (FT180100186). John McGrath is supported by the Danish National Research Foundation (Niels Bohr Professorship, the National Health and Medical Research Council (John Cade Fellowship 1056929). John McGrath, Darryl Eyles and Thomas Burne are employed by The Queensland Centre for Mental Health Research which receives core funding from the Queensland Health. Darryl Eyles is supported by the NHMRC (1124724, 1124721, 1141699). Brittany Mitchell received financial support from the Queensland University of Technology.

## Methods

### The UK Biobank sample

The UK Biobank (UKB) is a large population cohort with phenotype, genotype and clinical information on more than 502,000 individuals (age range from 40 to 69 years old). Participants were registered with the National Health Service, and lived approximately 25 miles from one of the 22 recruitment centres across the United Kingdom (UK)^11^. Participants were recruited between 2006 and 2010. Informed consent was obtained by UK Biobank from all participants, and the study was approved by the North West Multicentre Research Ethnics Service Committee. The participants of the study were not representative of the original sampling frame, with evidence of a ‘healthy volunteer’ bias^66^.

Genotype data were quality controlled and imputed to the Haplotype Reference Consortium (HRC)^67^ and UK10K^68^ reference panels by the UKB group^69^. We extracted variants with minor allele count (MAC) > 5 and imputation score > 0.3 for all individuals, and converted genotype probabilities to hard-call genotypes using PLINK2 (--hard-call 0.1)^70^. Then, we excluded variants with genotype missingness > 0.05, Hardy-Weinberg equilibrium *test P* > 1 × 10^−5^, and minor allele frequency (MAF) < 0.01. In total 8,806,780 variants (hereafter SNPs, but could include small insertion/deletions (INDELS)), including 260,713 SNPs in the X chromosome, were available for analysis.

Individuals of European ancestry were identified by projecting the UKB sample to the first two principal components (PCs) of the 1000 Genome Project (1KGP^71^), using Hap Map 3 (HM3) SNPs with MAF > 0.01 in both datasets. European ancestry was assigned based on > 0.9 posterior-probability of belonging to the 1KGP European reference cluster.

#### Assessment of 25 hydroxyvitamin D concentration

Vitamin D 25OHD levels were measured in blood samples collected at two instances: the initial assessment visit, conducted between 2006 and 2010, and a repeat assessment visit, conducted between 2012 and 2013. The Diasorin Liason®, a chemiluminescent immunoassay (CLIA) was used for the quantitative determination of 25OHD. The assay measures total 25OHD concentration (i.e. 25OHD_3_ and 25OHD_2_). Participants with 25OHD concentrations below or above the validated range for the assay (10 - 375 nmol/L) were excluded. The average within-laboratory coefficient of variation (CV) (and standard deviation) ranged from 5.04 (4.73) to 6.14 (2.21)^72^.

Of 502,536 UKB participants, 449,978 (90%) had vitamin D 25OHD levels (data field 30890) measured, mostly from the initial assessment visit (448,376, 99.6%). Our analyses were limited to the 417,580 individuals of European ancestry with 25OHD concentrations available, of whom 318,851 are unrelated (gcta --rel-cut-off 0.05).

#### Genome-wide association study (GWAS) analysis

**Figure 1** provides a graphical summary of the GWAS and post-GWAS analyses detailed below. To identify genetic variants associated with 25OHD levels, we performed a GWAS with fastGWA^73^. fastGWA is a tool implemented in GCTA^74^ for mixed linear model (MLM)-based GWAS. It uses a sparse genomic relationship matrix (GRM) to account for genetic structure within the cohort, making it a resource-efficient method for the analysis of large datasets like the UK Biobank^73^. The sparse GRM was generated for UKB individuals of European ancestry using HapMap3 SNPs.

We applied a rank-based inverse-normal transformation (RINT) to the phenotype (vitamin D 25OHD levels) and fit age at time of assessment, sex, assessment month, assessment centre, supplement intake information, genotyping batch and the first 40 ancestry PCs as covariates in the model (see **Supplementary Methods** for more details).

To identify independent associations, we a conducted a conditional and joint (COJO; gcta --cojo-slct) analysis^20^ of the GWAS results, accounting for the correlation structure between SNPs within a 10-Mb window and using a random subset of 20,000 unrelated Europeans from the UKB as linkage disequilibrium (LD) reference. For comparison, we used PLINK1.9 (--clump)^75^ to identify regional lead SNPs for genome-wide significant index variants (--clump-p1 5e-8) and variants were clumped with this lead SNP if they were located less than 10,000 Kb (--clump-kb 10000) away from, and with *r*^2^ > 0.01 (-- clump-r2 0.01) with the index variant. To identify novel associations, we a conducted a COJO analysis that conditioned (gcta --cojo-cond) on the six loci previously reported as genome-wide significant (rs2282679, rs10741657, rs12785878, rs10745742, rs8018720, rs6013897)^10,22,76^.

#### Meta-analysis

The largest GWAS for 25OHD to date, from the SUNLIGHT consortium^10^, used BMI as a covariate, hence we also generated UKB results including BMI in the model and used those for meta-analysis. In addition, the UKB GWAS results used for meta-analysis differed from the reported GWAS results in that 25OHD levels were natural-log transformed and supplement intake was not included as a covariate. Before meta-analysis, we imputed the SUNLIGHT summary statistics (2,579,297 SNPs) with ImpG^24^. After data management, we used a sample size-based approach^77^ to perform the meta-analysis (**Supplementary Methods**) on 6,912,294 SNPs that were shared between the data sets.

#### Relationship between vitamin D and Body Mass Index traits

High BMI is associated with lower concentrations of 25OHD^12^. For this reason, previous GWAS of 25OHD have included BMI as covariate in their analyses^10^. However, given that BMI is a highly heritable trait, covariate adjustment can induce collider bias^17^ and affect downstream analyses. To better understand the relationship between 25OHD and BMI we estimated the phenotypic and genetic correlation between them and used generalised summary-data-based Mendelian randomisation (GSMR)^18^ to test for statistical evidence for putative causal effects between the two traits. SNP instruments were selected with the default settings of the built-in GSMR clumping step. In addition, we conducted a multi-trait conditional and joint (mtCOJO) analysis^18^ to condition the 25OHD GWAS results on BMI GWAS summary statistics generated with the UKB^19^, an approach that was shown in simulations to be robust to induced collider bias when conditioning on a correlated trait^18^. A random subset of 20,000 unrelated individuals of European ancestry from the UKB was used as LD reference in the mtCOJO analysis.

#### Heritability and SNP-based heritability

Our UKB sample included a set of 58,738 individuals who were related with coefficient of relationship (*r*) > 0.2 to at least one other person in the set (“all relatives”). Among these, there was a set including all pairs with 0.4 < *r* < 0.6 (1^st^ degree), and set a including all pairs with 0.2 < *r* < 0.3 (2^nd^ degree). We used these sets to estimate heritability of RINT(25OHD) levels (gcta --reml). To estimate SNP-based heritability we drew a random subset (N ∼ 50,000), selected so that no pair of individuals had *r* > 0.05. We used a model that fits a single random genetic effect with a single genomic relationship matrix (GRM) constructed from all SNPs^78^ and also a GREML-LDMS model^79^ that fits 10 random genetic effects and hence 10 GRM (gcta --reml --mgrm). The 10 GRM were constructed from SNPs annotated to 5 MAF (0.01-0.1, 0.1-0.2, 0.2-0.3, 0.3-0.4, 0.4-0.5) bins each divided into two by median LD score of the SNPs within the bin. The LD score of a SNP is a measure of the common genetic variation tagged by a SNP. The sum of the estimates for each MAF-LD bin is an estimate of the total SNP-based heritability. Under a neutral model each of the 5 MAF bins is expected to explain 20% of the variance. Analyses were conducted with and without BMI as a covariate, and genetic correlation between 25OHD and BMI was estimated in a bivariate GREML analysis (gcta --reml-bivar). In addition, we estimated the genetic correlation and the genetic variance explained by 25OHD levels assessed in summer and winter (see definitions in ‘vQTL and seasonal analysis’ section below), using bivariate GREML. Heritability and SNP-based heritability estimated as part of the GWAS analysis using fastGWA are also reported. Finally, we estimated SNP-based heritability by LD score regression^32^ (software default settings for European ancestry samples), SBayesR^15^ and SBayesS^15^ from GWAS summary statistics. From SBayesS we also estimate the polygenicity and selection parameters.

### Replication and out-of-sample genetic risk prediction

We used the Brisbane-based twin and family sample (N = 6,223)^8^ for replication analyses. Samples were collected between May 1992 and January 2014, mostly from South-East Queensland (latitude 27° S). At this latitude, there is sufficient UVR to allow for vitamin D synthesis throughout the year^80^. Legal guardians gave written, informed consent prior to inclusion and testing. Studies were approved by the Human Research Ethics Committee of the QIMR Berghofer Medical Research Institute. Additional details of this study are provided elsewhere (25OHD assay methods^8^ and genotyping on HumanCoreExome-12v1-0_C or IlluminaHuman610W-Quad bead chip and quality control^81^). Genotypes were imputed to phase 3 version 5 of the 1000 Genomes Build37 (hg19)^23^. The phenotype analysed was RINT(25OHD) pre-regressed on sex, age, month of collection, 10 ancestry PCs and imputation batch. We selected a set of 1,632 unrelated individuals from the Brisbane cohort and estimated the proportion of variance explained by the UKB genome-wide significant SNPs (or their proxies) when fitted jointly by using only these SNPs to construct a GRM and conducting a GREML analysis in GCTA. Next, using the sample we conducted polygenic risk score (PRS) analyses. Using only SNPs present in the Brisbane cohort we selected independently associated SNPs from the UKB cohort in order to conduct standard *P*-value thresholding PRS analysis, choosing a range of *P*-value thresholds (*P* < 5 × 10^−8^, *P* < 1 × 10^−5^, *P* < 0.001, *P* < 0.01, *P* < 0.05, *P* < 0.1, *P* < 0.5, *P* < 1) and calculating PRS for each individual in the Brisbane cohort. We also calculated PRS from SNP weights estimated by COJO^82^, SBayesR^26^ and SBayesS^15^. The Bayesian methods better account for the complex relationship between strength of association of, and correlation between, SNP effect sizes. For each set of PRS we estimated the proportion of variance explained by the PRS in the Brisbane cohort using GCTA GREML to account for the family structure.

### Functional mapping and annotation of GWAS

We conducted a number of analyses to annotate the 25OHD GWAS results. First, we used the FUMA online pipeline^27^ to obtain gene-based, gene-set and tissue-specific annotations. Second, we used functional annotations provided in the LDSC software to partition SNP-based heritability into 53 functional categories^29^. Annotations included elements such as UCSC, UTRs, promoter and intronic regions, conserved regions and functional genomic annotations constructed using ENCODE^83^ and Roadmap Epigenomics Consortium data^84^. Third, we assessed the SNP-based heritability enrichment associated with different cell-types. Specifically, we applied LDSC analysis to the GWAS summary statistics using scores associated with cell-type-specific expression (as provided in the LDSC software)^85^.

To help prioritize putative causal genes with expression underlying 25OHD levels, we used the summary data-based Mendelian randomization method (SMR)^30^. SMR integrates GWAS and eQTL (expression quantitative trait loci, SNPs associated with gene expression) results with the aim of identifying pleiotropic or causal associations between a trait of interest and gene expression. We used eQTLs derived by the eQTLGen consortium from gene expression in whole blood^31^, using the largest sample, to date, for blood eQTLs (N = 31,684), identifying 15,504 genome-wide significant eQTLs. In general, SNPs controlling variation in one tissue are found to control variation in other tissues^86^, hence using the largest eQTL data set is the most powerful. Moreover, blood is a relevant tissue for vitamin D-related gene transcription^87^. Other relevant tissues are liver, skin and, given our hypotheses about the relationship between 25OHD and psychiatric disorders, the brain. To capture tissue-specific eQTLs in these tissues we used GTEX eQTL data sets, despite the fact that these data sets are much smaller than the eQTLGen sample (sun-exposed skin N = 369; non-sun-exposed skin N = 335; liver N = 153; sixteen brain regions N between 80 – 154). In addition, we used eQTLs identified in pre-frontal cortex (N = 1,866) from the PsychENCODE project^88^, and foetal brain samples (N = 120) from O’Brien et al.^89^ SMR significant results were declared at *P* < 0.05 / M per tissue, where M is the number SMR tests performed (i.e. the number of gene probes tested for the tissue, **Supplementary Table 13**). While significant SMR test results implicate a role for the eQTL gene, SNPs passing the conservative SMR heterogeneity in dependent instruments (HEIDI) test (*P*_HEIDI_ > 0.05) have robust support for the direct causal or pleiotropic relationships of the trait-associated SNPs influencing gene expression.

### Genetic correlations and putative causal relationships with other traits

Published epidemiological studies have provided an extensive set of hypotheses about causal relationships between vitamin D and a range of phenotypes^90^, including psychiatric and brain-related disorders^91^. To characterize the relationship between vitamin D and psychiatric traits, we conducted two sets of analyses. First, we used bivariate LD score regression^32^ to estimate the genetic correlation between vitamin D and psychiatric traits using the GWAS summary statistics generated with the UKB dataset and GWAS summary statistics that were publicly available for attention deficit/hyperactivity disorder (ADHD)^92^, Alzheimer’s disease (AD)^93^, major depressive disorder (MDD)^94^, schizophrenia (SCZ)^95^, bipolar disorder (BIP)^96^ and autism spectrum disorder (ASD)^97^. In addition, we obtained genetic correlation estimates between vitamin D and 746 traits available through the LD Hub database^34^. Second, we conducted generalized summary Mendelian randomization (GSMR) analyses^18^ to assess if there was any statistical evidence that observed correlations could be explained by a causal relationship for seventeen traits (**Figure 5**). GSMR analyses were conducted as described above (see BMI section), with significance declared at 0.05/18=0.003. For any significant associations observed with GSMR we confirmed our conclusions using the 2-sample MR (2SMR) method^33^, which implements a range of MR models that can adjust for the potential influence of pleiotropy (MR Egger, weighted mean, inverse variance weighted, simple mode, and weighted mode).

#### Proxy-environment vQTL and seasonal analysis

25OHD concentration is known to be affected by season of measurement, but other environmental factors may impact 25OHD measures. We conducted a genome-wide vQTL analysis^14^, an approach to detect presence of genotype-by-environment interaction in the absence of measurement or knowledge of the interacting environmental risk factor, to identify SNPs associated with variance in 25OHD. Specifically, we used the Levene’s median test implemented in OSCA^98^. Following the guidelines of Wang et al., we (1) adjusted 25OHD levels for selected covariates (see below), (2) removed outliers more than 5 SD from the mean, and (3) standardised the residuals to have mean 0 and variance 1. Each step was performed within one of eight groups defined based on sex (male vs. female) and supplement intake (none, other, vitamin D, or missing). This approach removed both the mean effect of covariates and the mean and variance differences between gender and supplement-intake groups, while retaining other distributional properties of the measure. Covariates included in the phenotype pre-regression were age at assessment, assessment month, assessment centre, genotyping batch and the first 40 PCs. To avoid spurious associations due to coincidence of low-frequency variants with phenotype outliers^14^, this analysis was restricted to SNPs with MAF > 0.05. To identify near-independent vQTLs, we clumped the vQTL GWAS results with PLINK1.9 (--clump) as above.

To assess if significant vQTL associations reflected a GxE with season of testing, we conducted season-stratified GWAS using PLINK1.9^70^(--gxe) and compared the results with the vQTL GWAS results. Specifically, we stratified the UKB cohort into two groups after visual inspection of the mean 25OHD concentrations per month (**Supplementary Figure 1b**). We defined two discrete time periods in order to retain the maximum sample size but optimize comparisons between months with higher and lower mean 25OHD concentrations: (a) “Winter” - individuals assessed Dec-Apr (N = 162,591), and (b) “Summer” - individuals assessed Jun-Oct (N = 177,082). Individuals with vitamin D levels assessed in the months of May and November were not included in these analyses.

## Data availability

Genome-wide association summary statistics generated with the three levels of BMI correction (i.e. with and without BMI as covariate, and conditioned on BMI) are available for download from http://cnsgenomics.com/data.html. Results for the UKB GWAS of BMI used for conditional analysis are also available from the same website.

## Author Contributions

JAR, JJMcG and NRW conceived the study and designed the analyses. JAR, TL, JQ, and BM conducted the analyses, KEK, and JS performed the initial preparation and quality control of the UK Biobank data. AX,YH,ZZ,JZ,HW,AAEV,GZ provided support in analysis implementation. JF,DE,THJB helped with interpretation of identified loci. NGM provided the QIMR cohort, and BM and GZ conducted the analyses based on this sample. PMV and JY provided advice on analyses and interpretation of results. JAR, JJMcG and NRW wrote the manuscript with the participation of all authors. All authors reviewed and approved the final manuscript

