## SUPPLEMENTARY MATERIAL for "Genome-wide association study identifies 143 loci associated with 25 hydroxyvitamin D concentration"

***Revez et al***

Supplementary Methods:	2

Supplementary Results:	3

Supplementary Figures:	4

Figure S1: Distribution of 250HD concentrations in the UK Biobank sample included in this study (N = 417,580)

Figure S2. Heritability and SNP-based heritability estimates of 25OHD levels estimated with different methods.

Figure S3: GREML-LDMS estimates of SNP-based heritability

Figure S4: Number of COJO GWS loci from random draws of samples from the UKB sample

Figure S5: Example SMR example plot

Figure S6: Genetic correlations estimated from bivariate LDSC regression using GWAS summary statistics from 25OHD and some selected phenotypes, using GWAS for 25OHD based on three types of adjustment for body mass index (BMI).

Figure S7: Distribution of -log_10_ *P*-values from 20 vQTLs without GWS GxE association with season of blood draw.

**Supplementary Methods**

*Supplement intake groups*

Supplement intake information was obtained from two UKB data fields – 104670 (indicating whether individuals replied to the survey on supplement intake) and 20084 (summarising which supplements were taken). Fields 104670 and 20084 were assessed at 5 instances. Using information from all timepoints, four groups were defined, namely: (1) none (individuals who always reported not taking supplements; N = 90,992), (2) other (individuals who ever reported taking other supplements, but never vitamin D; N = 76,863), (3) vitamin D (individuals who ever reported taking vitamin D, and may also have taken other supplements; N = 11,635), and (4) missing (individuals with no information on supplement intake in any of the five timepoints; N = 238,090).

*Principal components*

Principal components (PCs) were calculated with UKB genotype calls, which were LD pruned (LD r^2^ < 0.01) and had long-range LD regions removed by the UKB team. A total of 137,102 variants with MAF > 0.01, HWE *P-*value < 10^-6^ and genotype missingness < 0.05 in unrelated Europeans were used to calculate the PCs with flashPCA^1^. PCs were calculated for unrelated Europeans and projected onto the complete set Europeans.

*SUNLIGHT GWAS summary statistics*

The SUNLIGHT data were downloaded from <https://drive.google.com/drive/folders/0BzYDtCo_doHJRFRKR0ltZHZWZjQ> (file 25HydroxyVitaminD_QC.METAANALYSIS1.txt). In total, 79,366 individuals from 31 GWAS cohorts of European descent in Europe, Canada and USA were included in that study. The SUNLIGHT consortium performed genome-wide analyses within each cohort using a uniform analysis plan. Specifically, additive genetic models were fitted using linear regression on natural-log transformed 25OHD. Month of sample collection, sex, age, body mass index, and principal components (PCs) were included as covariates. Then, a fixed-effect inverse variance weighted meta-analysis was conducted using the METAL^2^ software package, with control for the population structure. SNPs with a MAF ≤ 0.05, imputation info score ≤ 0.8, HWE ≤ 1× 10^−6^, and less than two studies or 10,000 individuals contributing to each reported SNP association were removed. After quality control, the SUNLIGHT consortium made summary statistics on 2,579,297 SNPs available for download.

*Imputation of the SUNLIGHT summary statistics to 1KGP using ImpG*

Since individual-level genotypes are not available from the SUNLIGHT consortium, we imputed the SUNLIGHT summary statistics to 1000 Genome Project (1KGP) using ImpG^3^. The haplotype reference panel files (EUR) and SNP mapping files were obtained from 1KGP phase 1 (release v3). Since information on allele frequency was not available in the SUNLIGHT summary statistics, we imputed the allele frequency info using the 1000G imputed ARIC data. We removed SNPs that had mismatched alleles in the two cohorts. After imputation, we further removed SNPs with imputation accuracy metric (r2pred score) < 0.6, and were left with 7,667,712 SNPs. We extracted 6,912,294 SNPs that were available both in the imputed SUNLIGHT and the UKB summary statistics. We further checked the consistency of allele frequencies and the heterogeneity of effect sizes of the SNPs in both cohorts.

*Sample size based meta-analysis*

Since the effect size estimates (betas) and their standard errors (se) are not in entirely consistent units across the two cohorts, we adopted a sample size-based approach^2^ to perform the meta-analysis. This approach converts the direction of effect and *P*-value reported in each cohort into signed z-scores, which are combined across studies into a weighted sum (based on sample size) for each allele. However, since we have SNPs with very small *P*-values, which are recognized as zero by the METAL^2^ software and cannot be converted to z-score, we computed the z-score using the betas and se, and computed the meta-analysis using the same equations and tests in R as implemented in the METAL package.

Among the 6,912,294 SNPs, 6,165 SNPs showed heterogeneity (HetPVal < 1 x 10^-3^). Since many of these SNPs were very significant (758 SNPs with *P* < 5 x 10^-8^), we kept them for downstream analysis. 15,154 SNPs reached the GWS threshold (*P* < 5 x 10^-8^), and 150 SNPs and 166 SNPs were identified as independent signals by the GCTA-COJO and LD clumping analysis.

**Supplementary Results**

**Summary-data-based Mendelian randomization** **(SMR) analyses**

To identify 25OHD SNP associations with statistical evidence consistent with a causal/pleiotropic association via gene expression we used summary-data-based Mendelian randomization (SMR)^4^ using the 15,504 gene probes with significant cis-eQTLs identified from whole blood eQTLGen data^5^. After Bonferroni correction, we found 112 significantly-associated gene expression probes (*P*_SMR_ < 3.2 x10^-6^, i.e., 0.05 / 𝑚, with 𝑚 = 15,504, being the total number of probes tested in SMR analysis; (**Supplementary Table 13;** [**Supplementary Data**](https://www.dropbox.com/sh/rhimyqqswxnn4wk/AACErpIc2DmrwXoOl1XD-rFva?dl=0)). These genes were in 30 broad regions on 16 chromosomes (4 regions on chromosomes 1 and 4; 3 regions on chromosome 11; 2 regions on chromosomes 2, 3, 14, and 19; and 1 region on chromosomes 6, 7, 8, 9, 12, 15, 17 and 20). Genes identified in these analyses included those that encode proteins of interest to vitamin D-related pathways, such as *CYP2R1*, *DHCR7*, *NADSYN1*, *DGAT2* and *HSD17B11*. SMR also confirmed genes of interest discussed for the COJO analyses including *SDR42E1*, *DGAT2* and *POU2F3*. Of the 112 loci identified by SMR, 24 passed the HEIDI test (*P* > 0.05) and have the strongest statistical evidence for a causal role for vitamin D levels (the HEIDI test conservatively excludes results where the gene expression colocalization may be more complex than via the same set of causal SNPs). Of interest, SMR identified a *CYP24A1* intron eQTL variant (rs912505) - this enzyme catalyses additional hydroxylation of 1,25OHD, which leads to the degradation of the active form on vitamin D. SMR also identified eQTL variants related to the expression of (a) *AKR1A1;* a member of the aldo-keto reductase family 1, (b) two hydroxysteroid-related enzymes; *HSD17B13* and *HSD3B7,* and (c) *HAL*; an enzyme expressed in the skin that generates UVB-absorbing molecules.

We then used SMR to explore eQTLs in other relevant tissues (sun-exposed and non-sun-exposed skin, liver, and sixteen brain regions) using data from the GTExV7^5^, PsychENCODE^6^ and foetal brain tissue^7^ (**Supplementary Table 13;** [**Supplementary Data**](https://www.dropbox.com/sh/rhimyqqswxnn4wk/AACErpIc2DmrwXoOl1XD-rFva?dl=0)**)**. Of interest, we found significant SMR associations in many tissues for *RP11-660L16.2I*, which is an antisense non-coding RNA located in the bi-directional promoter between *DHCR7* and *NADSYN1*^8^. For both sun-exposed and non-sun-exposed skin, one of the top eQTL variants selected by SMR was rs3819817, within *HAL*. Overall, the SMR-related findings add weight to the hypothesis that these particular eQTL variants may be causally related to 25OHD concentrations. Plots for all significant SMR results are provided in the [**Supplementary Data**](https://www.dropbox.com/sh/rhimyqqswxnn4wk/AACErpIc2DmrwXoOl1XD-rFva?dl=0), with an example in **Supplementary Figure 5**.

**Supplementary Figures**

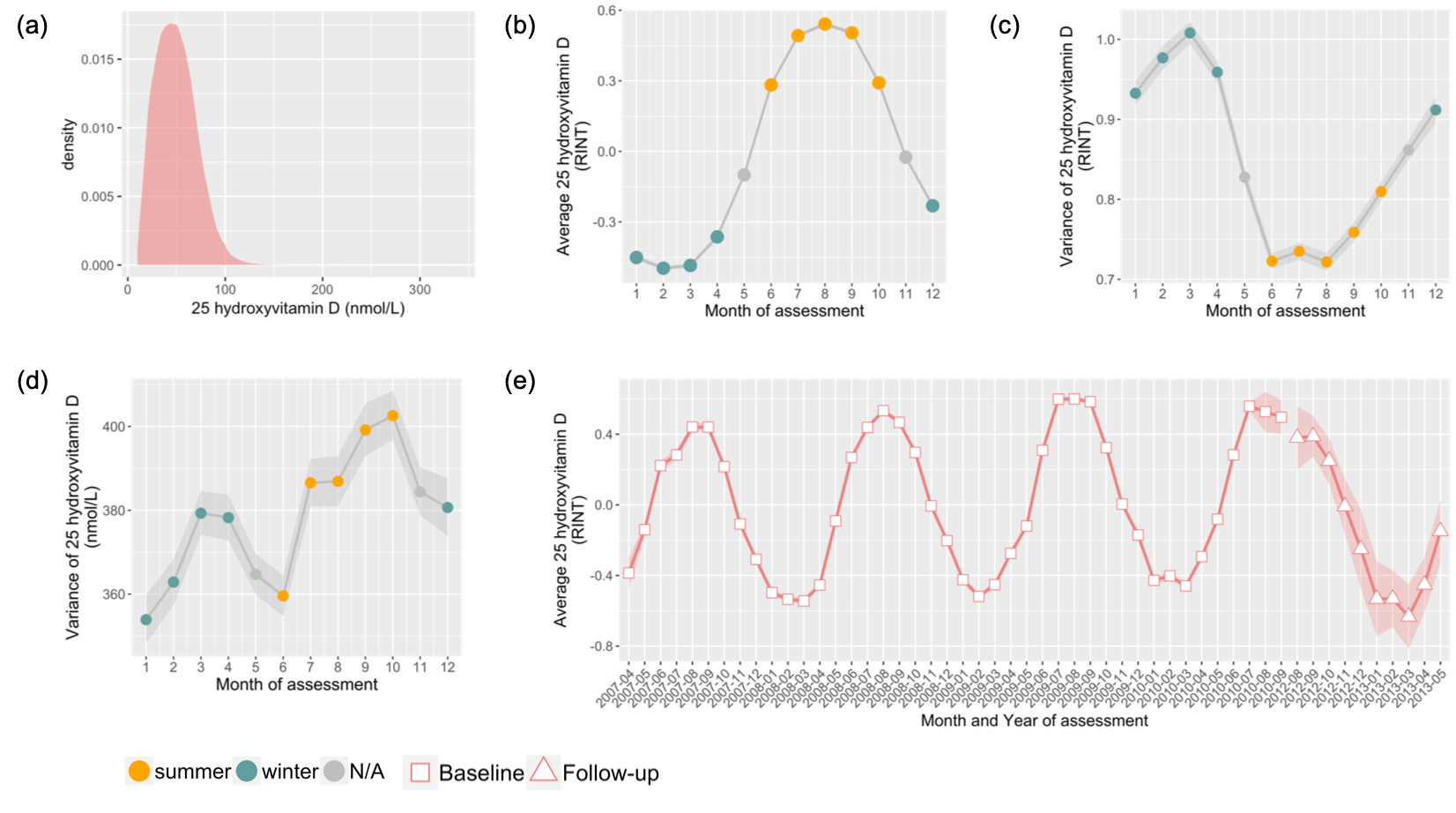

**Figure S1: Distribution of 25OHD concentrations in the UK Biobank sample included in this study (N = 417,580)**

**(a)** Density distribution of 25 hydroxyvitamin D (25OHD) concentrations (nmol/L). The right tail of the distribution had very low counts and is thus obscured by the grid lines. **(b)** Mean monthly 250HD concentrations after rank-based inverse-normal transformation (RINT). 95% confidence intervals are shown, but are obscured by the width of the line joining the monthly values. **(c)** Variance between individuals of monthly 25OHD after RINT. The lower variance in summer compared to winter is expected under a log transformation (because Var(log(x))~var(x) / mean(x)^2^), and also observed under RINT. **(d)** Variance between individuals of monthly 250HD (nmol/L). **(b-d)** Yellow and green dots represent months defined as summer and winter months, respectively. Grey dots represent months that were not included in season-stratified analyses. Shades represent 95% confidence intervals. **(e)** Mean monthly 250HD after RINT across years. Shades represent 95% confidence intervals. Squares represent measurements taken at the baseline visit. Triangles represent measurements collected from participants in first follow-up visit. Months with < 30 observations were not plotted.

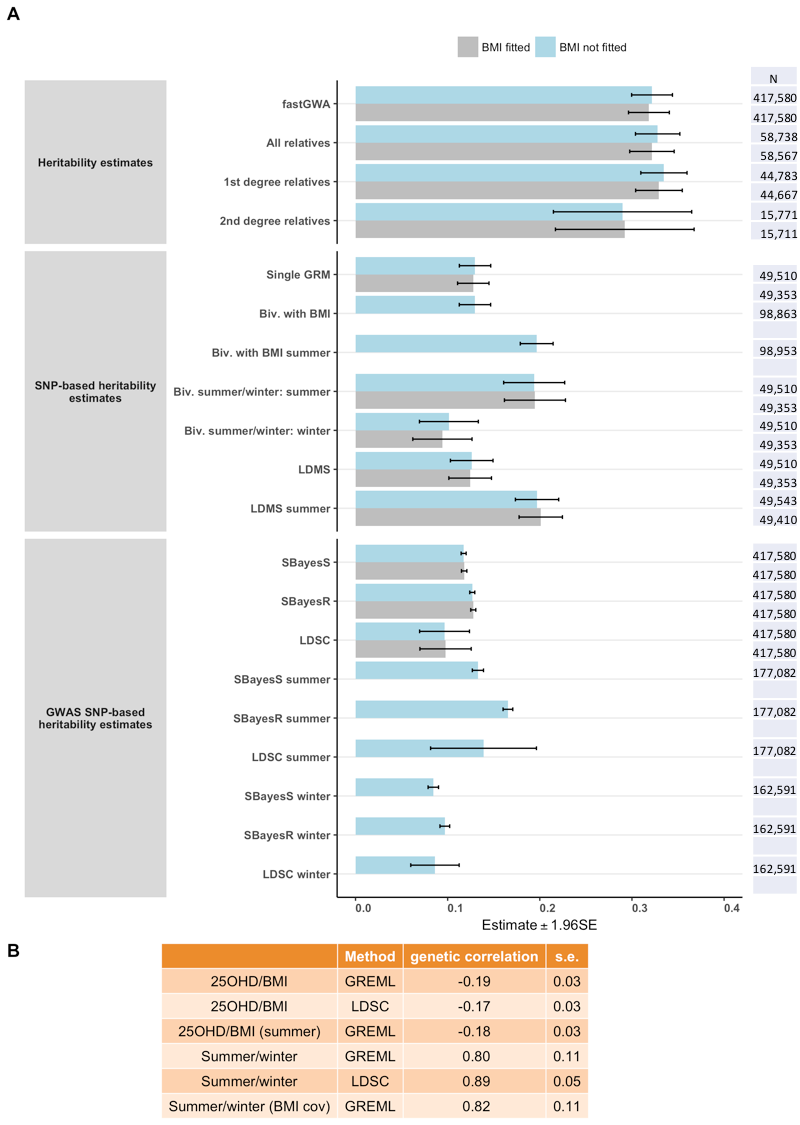

**Figure S2. Heritability and SNP-based heritability estimates of 25OHD levels estimated with different methods.**

Abbreviations: GRM, genomic relationship matrix; single GRM analysis contrasts to the linkage disequilibrium minor allele frequency stratified (LDMS) in which the linear mixed model has 10 genetic random effects with 10 GRM constructed from SNPs allocated to 10 bins based on their minor allele frequency and LD score annotation; Biv, bivariate; r_g_, genetic correlation.

Most methods were applied with (blue bars) and without (grey bars) BMI fitted as a covariate. Error bars represent the 95% confidence interval of the estimates. The sample size used in each analysis is listed on the right. All estimates are provided in **Supplementary Table 2**.

Our UKB sample included a set of 58,738 individuals related with coefficient of relationship (*r*) > 0.2 to at least one other person in the set (“all relatives”), from whom we estimate the heritability of 25OHD to be 0.32 (s.e. = 0.01). Among these, a set of 1^st^ degree relatives (pairs with 0.4 < *r* < 0.6; N = 44,783) and set of 2^nd^ degree relatives (0.2 < *r* < 0.3; N = 15,771) were identified. Heritability estimated from all relatives was 0.32 (s.e. = 0.01). Consistent with some confounding of genetic and shared environment factors between closer family relatives, the heritability estimated from 1^st^ degree relatives (0.33, s.e. = 0.01) was higher than that estimated from 2^nd^ degree relatives alone (0.29, s.e. = 0.04), but the difference was not significant (*P* = 0.33;

Bivariate GREML was conducted between 25OHD and BMI, but only the 25OHD heritability estimates are plotted. Genetic correlation (r_g_) estimates and respective standard errors (s.e.) were also obtained from the bivariate GREML analyses. Estimates of genetic correlation between 25OHD and BMI obtained with LDSC regression were -0.17 (s.e. = 0.03) using the UKB data and -0.12 (s.e. = 0.03) using the published BMI meta-analysed results, which excluded the UKB^9^. Bivariate GREML SNP-based (co)heritability analysis estimated a genetic correlation of -0.19 (s.e. = 0.03) and correlation of residual effects of -0.21 (s.e. = 0.05) with BMI.

We also estimated SNP-based heritability using three summary-data-based methods: SBayesR ^10^, SBayesS^11^, and LDSC^12^. As expected^13^, estimates from GWAS summary statistics were slightly lower than GREML estimates. SBayesR, a Bayesian method that models SNP effects based on a mixture of normal distributions and a point mass at zero, gave a SNP-based heritability estimate of 0.13 (s.e. = 1.4 x 10^-3^). SBayesS is another Bayesian method to estimate SNP-based heritability, which assumes a zero-normal mixture distribution for the SNP effect, with effect size-MAF relationship incorporated as a free parameter (S). In SBayesS analyses, the S parameter was -0.78 (s.e. = 0.04; **Supplementary Table 4**), consistent with a model of negative selection on the genetic variants associated with 25OHD levels (the magnitude of S is higher than those of most complex traits studied by Zeng at al.^11^). Heritability estimates obtained with LDSC (0.10, s.e. = 0.01) were comparable, but less accurate (larger standard errors), than those obtained with SBayesS. This is likely due to the more conservative method (Jackknife) that LDSC uses to estimate the variance of the estimate.

In summer, mean 25OHD levels were always higher than in winter (58.0 vs 41.2 in nmol/L; 0.4 vs. -0.4 RINT SD) (**Supplementary Table 1**). On the other hand, the standard deviation of 25OHD levels measured in summer was only higher than winter when assessment was made on the observed scale (19.8 vs 19.3 in nmol/L; 0.9 vs. 1 in units of RINT(25OHD) applied to the total sample; **Supplementary Figure 1c and 1d**). Regardless of the method used for the analysis, the SNP-based heritability for 25OHD levels assessed in summer was always higher than in winter. The genetic correlation between the seasons is 0.80 (s.e.= 0.11). While this is not significantly different from one, there may be genetic heterogeneity in 250HD levels between summer and winter, as investigated in GxE analysis.

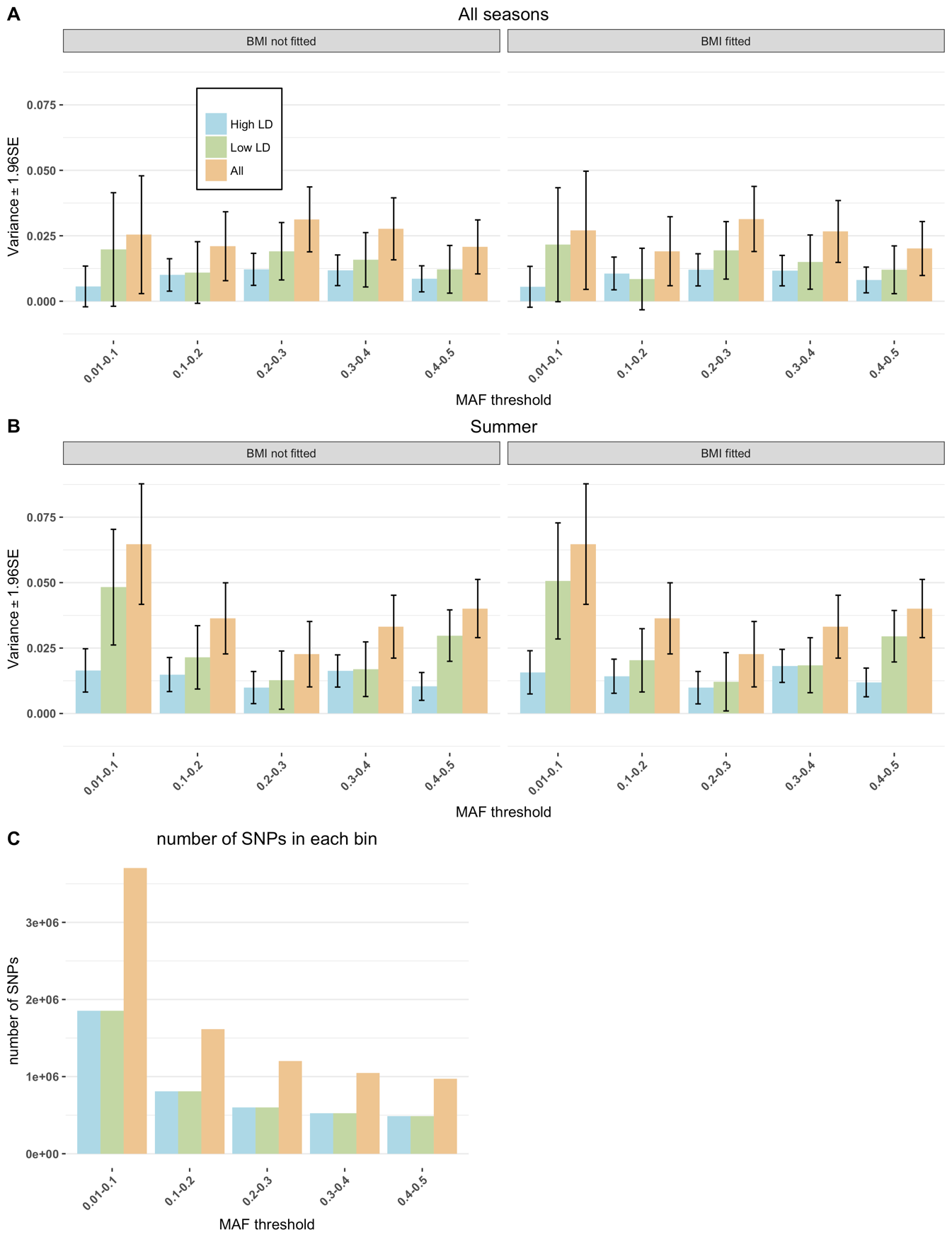

**Figure S3: GREML-LDMS estimates of SNP-based heritability**

(**A-B)** SNP-based heritability estimates stratified by five MAF groups (0.01 – 0.1; 0.1 – 0.2; 0.2 – 0.3; 0.3 – 0.4, 0.4 - 0.5) and two LD groups. Blue for High LD, green for Low LD, and orange for all NSPs in each MAF bin without LD stratification). **(A)** SNP-based heritability estimates from 25OHD assessments in all months of the year. **(B)** SNP-based heritability estimates from 25OHD assessed in summer months (June – October). (**C)** Number of SNPs in each of the 5 MAF x 2 LD bins.

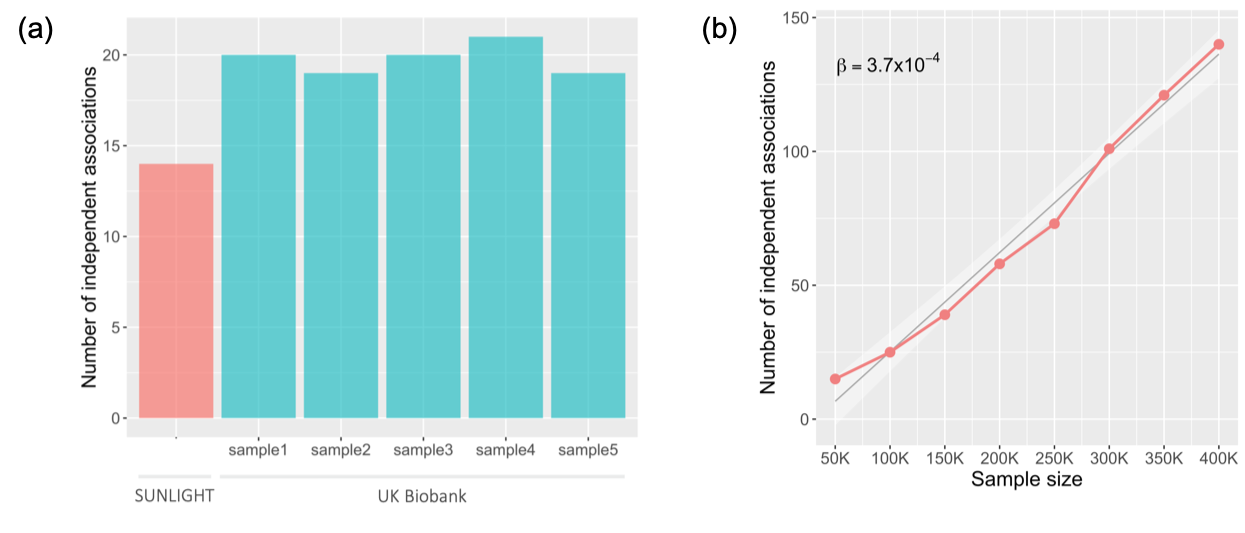

**Figure S4: Number of COJO GWS loci from random draws of samples from the UKB sample**

**(a)** Five random draws of 80K samples (the same size as the SUNLIGHT consortium) identify ~20 genome-wide significant (GWS) loci. SUNLIGHT identifies 14. **(b)** There is an approximately linear trend of 3.7 number loci per 10,000 individuals.

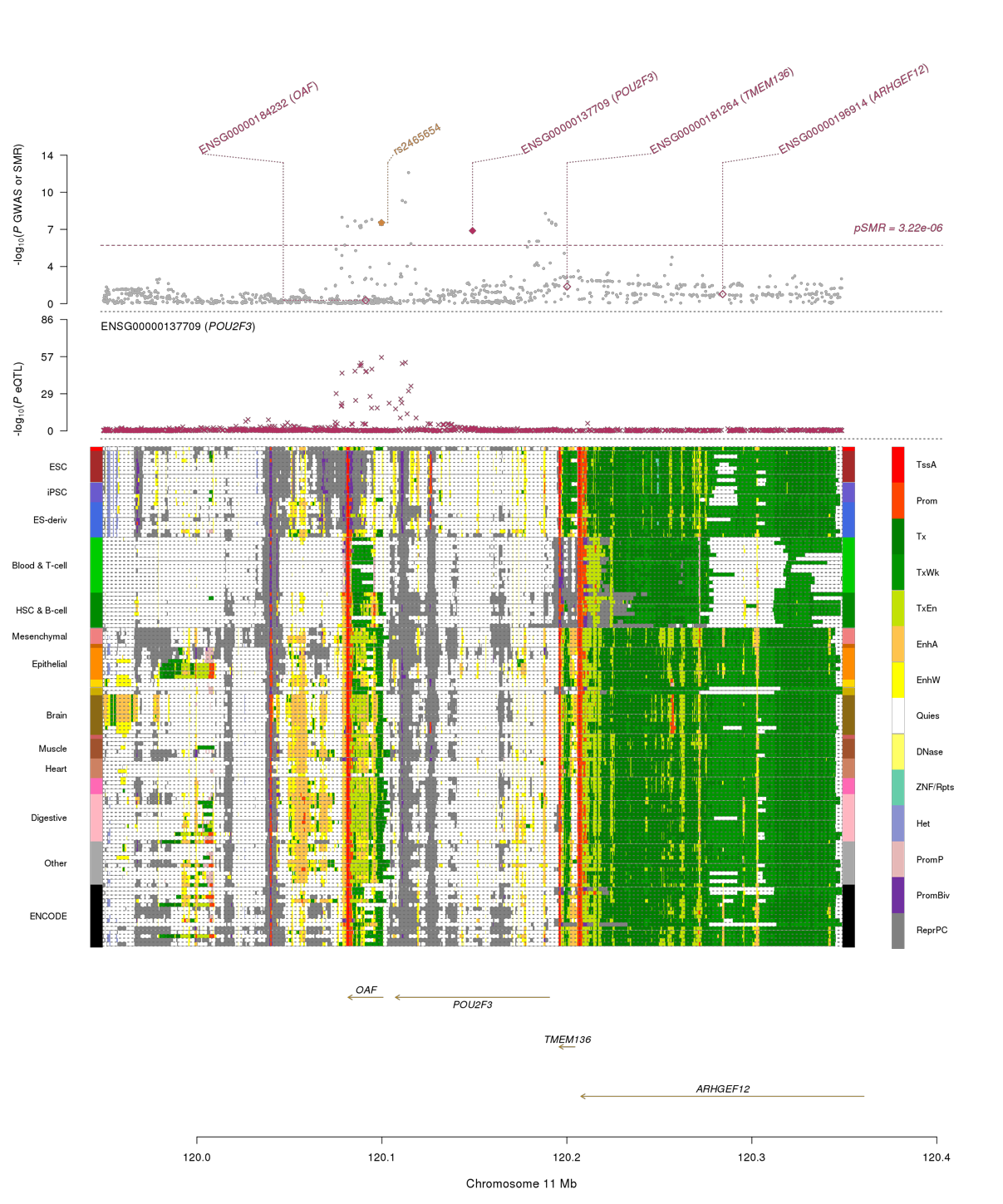

**Figure S5: Example SMR plot**

The above figure shows the results of the SMR+HEIDI analysis that combines the data from GWAS and eQTL studies. All results are provided in the [**Supplementary Data**](https://www.dropbox.com/sh/rhimyqqswxnn4wk/AACErpIc2DmrwXoOl1XD-rFva?dl=0) file. Amongst the eQTLs tested, rs2465654 is an eQTL for *POU2F3* in blood and liver (of tissues tested) and generates significant SMR associations to 25OHD. Notably, rs2465654 is not an eQTL in GTEX skin tissue, despite *POU2F3* having massively higher expression in skin compared to other tissues, and where it plays a critical role in keratinocyte proliferation and differentiation. The first track shows -log_10_(P-value) of SNPs (grey dots) from the GWAS of 25OHD. Each red rhombus indicates the -log_10_(P-value) from the SMR tests for associations of gene expression with 25OHD concentration. A solid rhombus represents a probe not rejected by the HEIDI filtering. The yellow rhombus denotes the SNP-25OHD association of the SNP that is the top cis-eQTL (rs2465654). The second track shows -log_10_(P-value) of the SNP association with gene expression probe ENSG00000137709 (tagging *POU2F3*). The third track displays information for 25 chromatin states (indicated by the colours on the right bar) of 127 samples from Roadmap Epigenomics Mapping Consortium (REMC)^14^ for different tissues and cell types (indicated by the colours on the left bar), which communicates potential tissue-specific roles for the SNP. The bottom track shows the genes underlying the genomic region.

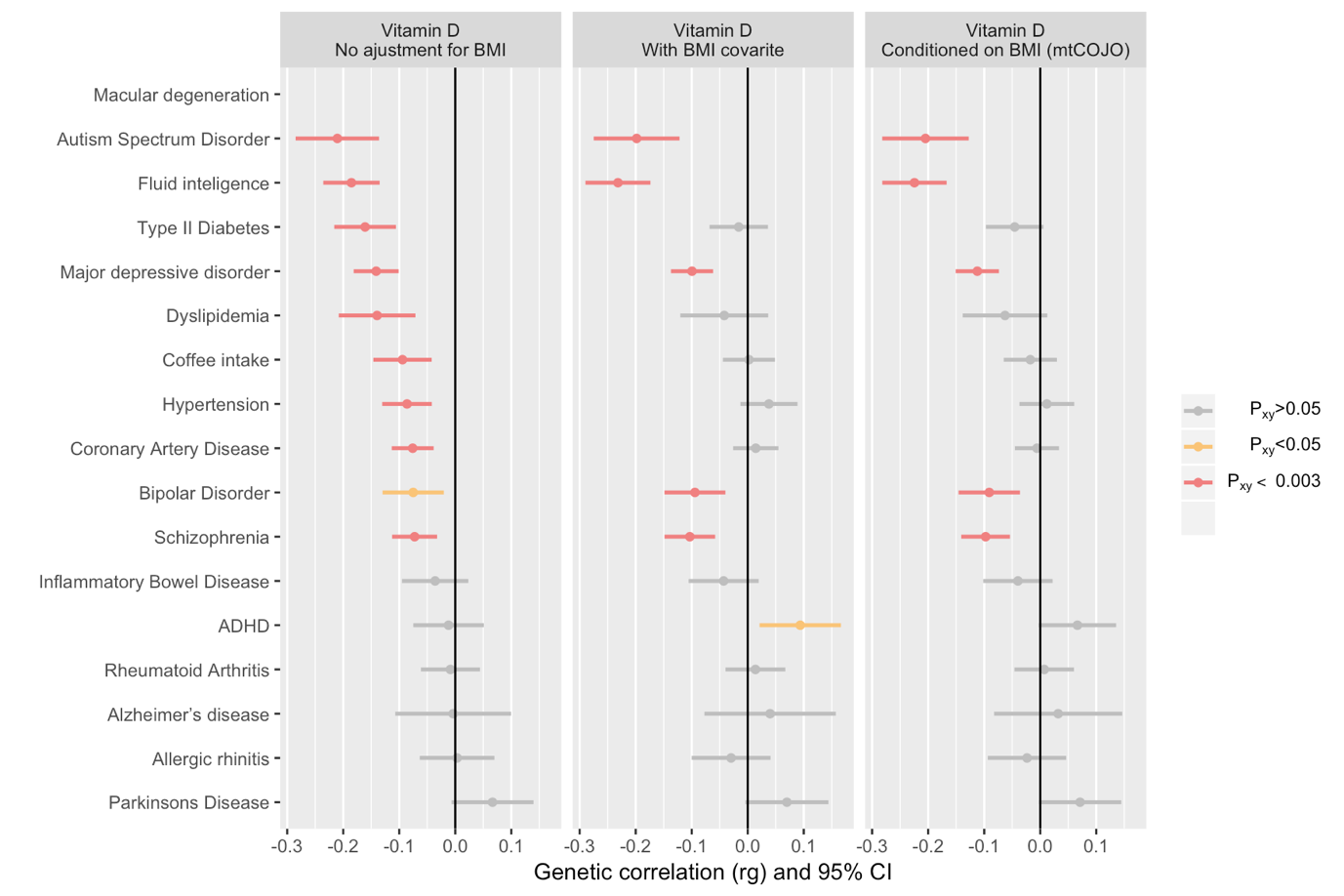

**Figure S6: Genetic correlations estimated from bivariate LDSC regression^12^ using GWAS summary statistics from 25OHD and some selected phenotypes, using GWAS for 25OHD based on three types of adjustment for body mass index (BMI).**

The genetic correlations and 95% confidence intervals between variants associated with 25 hydroxyvitamin D (25OHD) and selected phenotypes. Correlations are shown for analyses that make no adjustment for BMI, include BMI as a covariate, or condition on genetic correlates of BMI (via mtCOJO)^15^. Negative genetic correlations indicate the variants associated with increased 25OHD concentration were associated with a smaller value/reduced risk for the phenotypes of interest. The colours of the bars indicate *P-*values for each genetic correlation (grey *P* > 0.05, yellow 0.003 < *P* < 0.05, red *P* < 0.003).

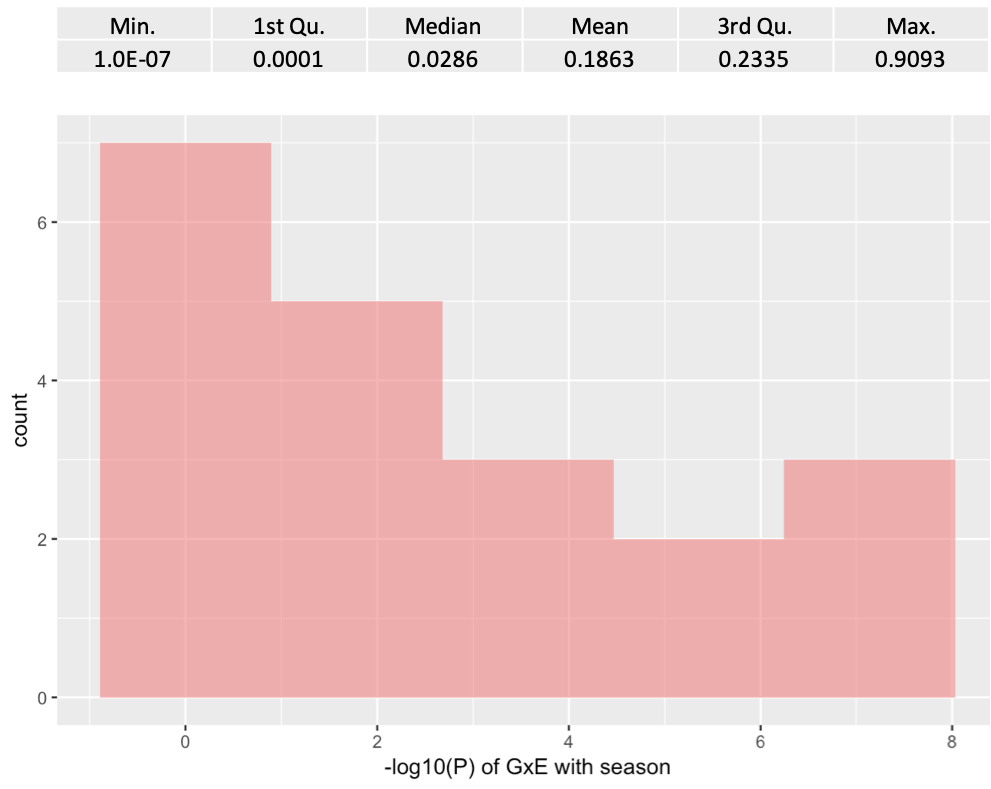

**Figure S7: Distribution of -log_10_ *P*-values from 20 vQTLs without GWS GxE association with season of blood draw.**

The median p-value is 0.03, which suggests that at least half of these 20 vQTL are unlikely to have SNP*season interaction.
